# CBL mutations in pediatric solid and CNS tumors are a marker of receptor tyrosine kinase activation and a potential therapeutic target

**DOI:** 10.64898/2025.12.02.691063

**Authors:** Lauren M. Brown, Chelsea Mayoh, Katherine A. Camper, Gabor Tax, Pablo Acera Mateos, Wenyan Li, Sam El-Kamand, Patricia Sullivan, James Bradley, Paris I. Thompson, Jingwei Chen, Teresa Sadras, Robert Salomon, Marie Wong, Mark J. Cowley, Antoine de Weck, Loretta M.S. Lau, Neevika Manoharan, Paul G. Ekert

**Affiliations:** Children’s Cancer Institute, Lowy Cancer Research Centre, UNSW Sydney, Sydney, NSW, Australia; School of Clinical Medicine, UNSW Sydney, Sydney, NSW, Australia; Kids Cancer Centre, Sydney Children’s Hospital, Randwick, NSW, Australia; Olivia Newton-John Cancer Research Institute, Heidelberg, Melbourne, Australia; School of Cancer Medicine, La Trobe University, Bundoora, Melbourne, Australia; Peter MacCallum Cancer Centre, Parkville, VIC, Australia; The Sir Peter MacCallum Department of Oncology, University of Melbourne, Parkville, VIC, Australia

## Abstract

Mutations in *CBL*, an E3 ubiquitin ligase that negatively regulates receptor tyrosine kinases (RTKs) and potentiates intracellular signaling, have been extensively characterized in hematological malignancies. However, the impacts are unknown in other cancer types. We have identified established and novel aberrations in *CBL* in molecularly diverse pediatric CNS and solid tumors. Additionally, we present novel *CBL* splice mutations, including in the germline of a high-grade glioma patient, and alternative *CBL* splicing without corresponding genomic alterations. Functionally, we demonstrate that CBL exon 8/9Δ, typically associated with acute myeloid leukemia, is RTK activating in neuroblastoma and that novel *CBL* variants, CBL E366_E373del and CBL C384G, can cooperate with overexpression of EGFR to transform Ba/F3 cells. Collectively, these data represent a novel form of RTK activation in pediatric patients with solid and CNS tumors who might benefit from RTK-targeted therapies.

**Statement of significance:** Molecularly targeted therapies improve childhood cancer outcomes. Therapies targeting genes called receptor tyrosine kinases are particularly effective. We have identified new genetic alterations that may be a marker for receptor tyrosine kinase activation and drug sensitivity, enabling more patients to benefit from targeted treatments.

## Introduction

Despite overall improvements in survival of children diagnosed with cancer in the last 30 years(1), outcomes for patients diagnosed with specific tumor types remain poor. Pediatric high-grade glioma (pHGG) is one such example, which accounts for a disproportionally high level of pediatric CNS tumor deaths (40%) despite its relatively low incidence (10%), and remains a clinically challenging disease to treat(2). Precision medicine programs, like the Zero Childhood Cancer Program (ZERO), are improving outcomes for children diagnosed with high-risk and difficult-to-treat cancers, like pHGG, by identifying molecular targets for therapy(3). Critically, ZERO has demonstrated that molecularly targeted therapies, or precision-guided treatments (PGTs), can improve survival in pediatric cancer patients with high-risk disease (2-year PFS: 26% vs. 5.2% for unguided treatment)(4). Further to this, comprehensive molecular analysis of pediatric cancer samples, as performed by ZERO, can enable the detection of novel cancer drivers and molecular targets(5), increasing the number of patients that can receive PGTs. Small molecule tyrosine kinase inhibitors (TKIs), targeting either non-receptor receptor tyrosine (NRTK) or receptor tyrosine kinase (RTK) genes, account for much of the success of molecularly targeted therapies in adult(6, 7) and paediatric cancers(8–10). Given this, the identification of molecular aberrations in NRTK and RTK genes is a high priority in precision medicine. It is clear, however, that we are likely missing opportunitites to use these effective drugs in more patients.

CBL Proto-oncogene, *CBL*, encodes a RING finger E3 ubiquitin ligase and signaling molecule that plays a key regulatory role in RTK and intracellular signaling(11). As an E3 ubiquitin ligase, CBL binds ubiquitously to activated (and phosphorylated) RTKs, e.g. EGFR, mediating their ubiquitination and subsequent degradation. As a signaling molecule, phosphorylated CBL binds to adaptor proteins, including SRC and CRKL, promoting activation of intracellular signaling pathways (e.g. PI3K/AKT) (12, 13). Oncogenic forms of CBL contain mutations in a critical alpha-helix region, located between the tyrosine kinase binding (TKB) and RING finger domains. This includes deletion of single tyrosine residues, ΔY368 and ΔY371, and larger intragenic deletions. These deletions include the 70Z Cbl isoform, which lacks 17 amino acids (E366_K382) in exon 8 and is the result of a splice acceptor site mutation, and the p95 Cbl isoform, lacking 111 amino acids (E366_K477) and corresponding to deletion of exon 8 and 9, all originally identified in mouse models(14, 15). These mutant forms of Cbl can block Cbl-mediated polyubiquitination of RTK targets and induce cellular transformation(16). In contrast, missense mutations affecting the same region, while still unable to mediate polyubiquitination of RTK targets, are unable to transform cells alone.

In human malignancies, *CBL* variants were first identified and characterized in acute myeloid leukemia (AML)(17, 18). These included missense variants, in the linker or RING finger domain, or splice region variants, resulting in the loss of exon 8, and occurred in the context of wild-type FLT3. Functionally, the CBL R420Q mutation was shown to cooperate with Flt3 to transform Il-3-dependent murine 32D cells(18). Studies on larger cohorts of patients with AML or myelodysplastic syndrome identified aberrant *CBL* isoforms lacking either exon 8 or exon 8/9, the result of either a segmental deletion or splice mutations, or missense mutations, which while rare (1-2%) associated specifically with core-binding factor and 11q deletion subtypes(19, 20). *CBL* mutations, including missense mutations and splice mutations, were subsequently shown to be associated with 11q acquired loss of uniparental disomy in myeloid malignancies(21–23). Studies in *c-Cbl^-/-^* hematopoietic stem/progenitor cells (HSPCs) showed that expression of *CBL* mutants increased responsiveness to cytokine stimulation, that was completely abolished when co-expressed with wild-type CBL, suggesting that this loss of wild-type CBL may be required(23).

In pediatric cancers, *CBL* mutations are most notably associated with Juvenile myelomonocytic leukemia (JMML)(24, 25). *CBL* mutations occur in approximately 15% of JMML patients, are associated with 11q uniparental disomy, and are predominantly of germline origin(24, 26, 27). Germline variants in *CBL* were subsequently associated with a developmental disorder, now characterized under the banner of RASopathies and referred to as CBL Syndrome(28), defined by Noonan syndrome-like phenotypic features and a predisposition to JMML(26). Germline *CBL* mutations include missense variants affecting the linker region and RING finger domain, most commonly affecting Y371, C381 and C384 residues, and splice site mutations, impacting either intron 7 (c.1096-1G) or intron 8 (c.1228–2A) splice acceptor sites resulting in partial or complete loss of exon 8 or 9(26). Germline and somatic variants in *CBL* have not been described for other pediatric cancers, outside of myeloid malignancies.

We have identified established and novel *CBL* variants in novel tumor contexts. Through the analysis of molecular data from patients enrolled in the ZERO program, we identified 26 individual *CBL* variants in 22 patients, which were enriched in patients with CNS tumors (12 patients), the majority of which were pHGG (7 patients). These variants included established CBL variants (CBL ex8/9Δ) in novel tumor contexts (neuroblastoma and germ cell tumors) as well as Variants of Uncertain Significance (VUS) and novel variants. We demonstrate that aberrant *CBL* splicing occurs in the context of novel splice region variants and in the absence of an identified genomic mutation. Functionally, we show that CBL ex8/9Δ can block CBL mediated degradation of activated RTK signaling and mediate resistance to TKIs, likely through activation of intracellular signaling through alternative pathways, in neuroblastoma cell lines. Further, characterization of novel variants identified in HGG patient samples in Ba/F3 cells demonstrated that these variants could cooperate with EGFR overexpression to drive cytokine independent proliferation. Together, this study shows that CBL mutations may be a marker of RTK activation and a therapeutic target in CNS and solid tumors. This finding will extend the benefit of TKIs, which have proven efficacy in pediatric cancer, to more patients for whom novel therapeutic strategies are urgently needed.

## Results

### CBL mutations occur in pediatric CNS and solid tumors

We obtained processed whole genome sequencing (WGS) and RNA sequencing (RNAseq) data from patients enrolled on ZERO that had identified mutations in *CBL*. Thirty-three somatic variants (in 31 individual samples from 28 patients) were identified and 3 germline variants (in 3 patients) (Fig. 1A and Supplementary Tables S1-S3). Most CBL variants were identified in non-hematological tumors, with the majority identified in CNS tumor samples (n=22), in addition to single cases in neuroblastoma, sarcoma and solid tumors (Fig. 1B). The majority of *CBL* variants (72% [26/36], including 3 germline and 23 somatic) clustered around key functional domains, the linker region and RING finger domain, forming a clear variant hotspot and were predicted to be pathogenic based on their location (Fig. 1A). In silico modelling of missense mutations identified (n=17 individual mutations) supported the predicted pathogenicity of variants in this region (Supplementary Fig. S1).

**Figure 1.**
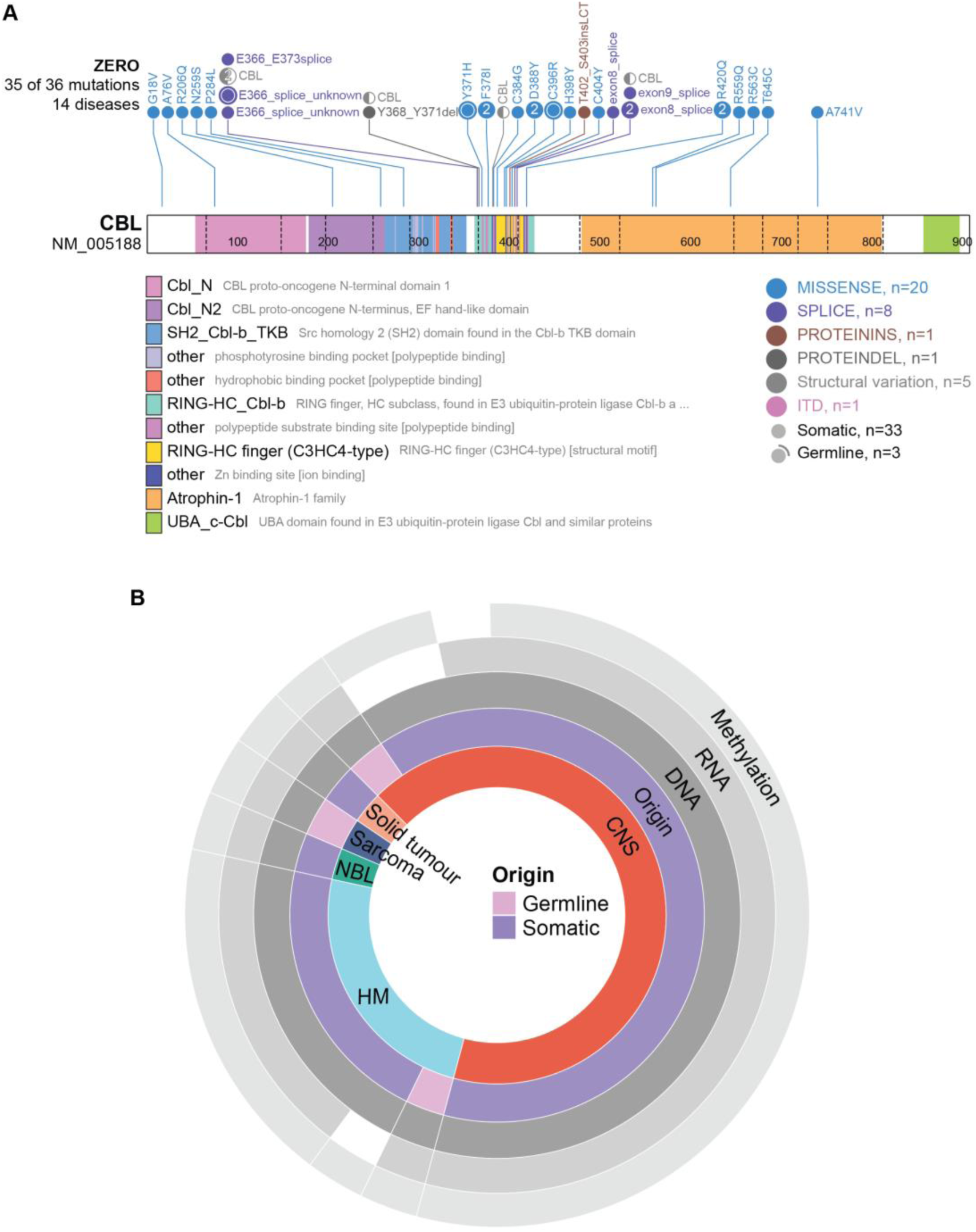
CBL mutations occur in pediatric solid and CNS tumors. A) Landscape of CBL mutations in the ZERO cohort (Protein Paint) including 10 predicted benign and 26 predicted pathogenic. B) Distribution of tumor types, CBL mutant origin (germline/somatic) and availability of molecular data from CBL-mutated samples.

Germline variants identified included an established pathogenic variant, CBL Y371H, in a juvenile myelocytic leukemia (JMML) (zccs1664) (26), a VUS, CBL C396R, in an embryonal rhabdomyosarcoma (RMS) (zccs932), and a novel splice region VUS (c.1096-3_1096-2del) in a HGG (zccs1267; Table 1 and Supplementary Table S4). Information about clinical features were available for the first two patients (zccs1664 and zccs932), for which they were concluded to have no clinical features of Noonan syndrome. This information was unavailable for the third patient (zccs1267). Of note, none of these patients had clinical features of Noonan syndrome. The identification of a pathogenic *CBL* germline variant, CBL Q367P, has been previously described in a single case of embryonal RMS with clinical features of Noonan syndrome(29). In addition, there is also a single case report of a VUS germline *CBL* variant, CBL L493F, in a patient with a H3K27M mutant diffuse midline glioma and a previous diagnosis of AML(30). Predicted functional somatic *CBL* variants were identified in 21 individual samples from 19 patients, including two patients with *CBL* variants identified at diagnosis and relapse (zccs444; HGG), and two stages of disease progression (zccs1262; CNS germ cell tumor).

**Table 1.** CBL mutation and clinical features of 22 paediatric cancer patients with likely functional *CBL* variants.

| ID | CBL variant | Significance | Germline | Tumour type | Stage |
| --- | --- | --- | --- | --- | --- |
| **zccs422** | p.F378I | VUS |  | HGG | D |
|  |  |  |  |  | R |
| **zccs1382** | p.C384G | Novel |  | HGG | D |
| **zccs1557** | p.R420Q | Likely Pathogenic |  | HGG | R |
| **zccs372** | p.R420Q | Likely Pathogenic |  | HGG | R |
| **zccs231** | c.1096-1G>T (splice) | Pathogenic |  | HGG | D |
| **zccs1267** | c.1096-3_1096-2del (splice) | Novel | + | HGG | D |
| **zccs1255** | c.1228-1G>A (splice) | Likely Pathogenic |  | HGG (radiation-induced) | R |
| **zccs2284** | c.1096-1G>T (splice) | Pathogenic |  | pilocytic astrocytoma | P |
| **zccs1528** | c.1096-1G>T (splice) | Pathogenic |  | pilocytic astrocytoma | P |
| **zccs2619** | c.1223_1227+10del (splice) | Novel |  | pilocytic astrocytoma | P2 |
| **zccs1468** | p.D388Y | Novel |  | CNS germ cell tumour | R1 |
|  |  |  |  |  | R2 |
| **zccs1704** | p.C404Y | VUS |  | ganglioglioma | P |
| **zccs1351** | p.exon8/9Δ | Pathogenic |  | neuroblastoma | R |
| **zccs1328** | p.exon8/9Δ | Pathogenic |  | malignant germ cell tumour | R |
| **zccs583** | p.exon8/9Δ | Pathogenic |  | AML | R |
| **zccs1847** | p.exon9Δ | Likely Pathogenic |  | AML | D |
| **zccs2395** | p.Y368_Y371del | Novel |  | AML | D |
| **zccs1103** | p.H398Y | VUS |  | AML | P |
|  | p.T402_S403insLCT | Novel |  |  |  |
| **zccs308** | c.1227+3_1227+7del ACGGA (splice) | Novel |  | AML | D |
|  | c.1227+8T>C (splice) | Novel |  |  |  |
| **zccs2223** | p.eK382_K477del | VUS |  | B-ALL | D |
| **zccs1664** | p.Y371H | Pathogenic | + | JMML | P |
| **zccs932** | p.C396R | VUS | + | Embryonal RMS | D |
| VUS: variant of uncertain significance, PA: pilocytic astrocytoma, HGG: high-grade glioma, AML: acute myeloid leukaemia, RMS: rhabdomyosarcoma, P: progression, D: diagnosis, R: relapse |  |  |  |  |  |

Somatic *CBL* variants were identified in 6 leukemia samples, including 5 AML and one B-ALL. These included two AML samples harboring established genomic deletions of exon 8-9 (CBL ex8/9Δ) or sub clonal deletion of exon 9 (CBL ex9Δ) and three AML samples harboring novel/VUS CBL missense mutations, splice mutations or indels. Partial deletion of exon 8 and exon 9 (K382-K477del), including part of the RING finger domain, was also identified in a B-ALL sample. The remaining samples harboring *CBL* variants (73% [16/22]) were from patients with non-hematological malignancies, including one neuroblastoma, one germ cell tumor, and 14 CNS tumor samples. The neuroblastoma (zccs1128) and germ cell tumor (zccs1253) samples both harbored CBL ex8/9Δ, representing the first identification of this mutation in these tumor types. The 14 CBL-mutated CNS samples included 3 pilocytic astrocytoma, 2 CNS germ cell tumor (matched progression samples from one patient), 1 ganglioglioma, and 8 HGG (from 7 patients). CNS tumors harbored *CBL* missense (8 samples) or splice (6 samples) mutations that were either novel, VUS and/or not previously identified in this tumor type. We conclude that the majority of pathogenic CBL variants in our cohort are occurring in CNS, as opposed to hematological cancers.

### *CBL* mutations occur in molecularly diverse tumors with features of receptor tyrosine kinase activation

Given the comprehensive molecular profiling performed by ZERO, we interrogated WGS, RNAseq, and methylation data from tumors with predicted pathogenic *CBL* mutations. As expected, *FLT3* aberrations were associated with *CBL* mutation in AML, with 60% of cases (3/5) harboring either *FLT3* mutation or elevated RNA expression (Fig. 2). Of the 16 solid and CNS tumor samples with somatic *CBL* variants, 63% of samples (10/16) harbored reportable (defined by ZERO as either pathogenic or likely pathogenic) genomic or transcriptomic events involving RTK pathway genes. These included genomic lesions associated with the disease, for example, *KIAA1549::BRAF* gene fusions in 3 patients with pilocytic astrocytoma and *KIT* missense mutations, identified at both timepoints, in a patient with a CNS germ cell tumor. These samples generally classified to DNA methylation-based subtypes associated with these mutations (80% [4/5 samples]). The remaining samples had aberrant RNA expression of intracellular kinase genes, notably *MTOR*, *HCK* and *MAPK* pathway genes (n=4) or mutation of *PTPN11* (n=1). Analysis of RNAseq data from *CBL*-mutated samples against the ZERO cohort showed that even within tumor types, *CBL*-mutated samples have distinct gene expression profiles (Supplementary Fig. S2). Further, *CBL*-mutated CNS tumor samples had distinct DNA methylation profiles and commonly did not classify to pre-defined DNA methylation-based subtypes (CNS v12.5(31)), which is in contrast to other CNS tumours from ZERO that more frequently classified(3) (Supplementary Fig. S3 and Supplementary Table S4). Interestingly, 2 HGG samples (zccs1382 and zccs422(D)) classified to subtypes associated with RTK activation, HGG_E and pedHGG_RTK1A, respectively.

**Figure 2.**
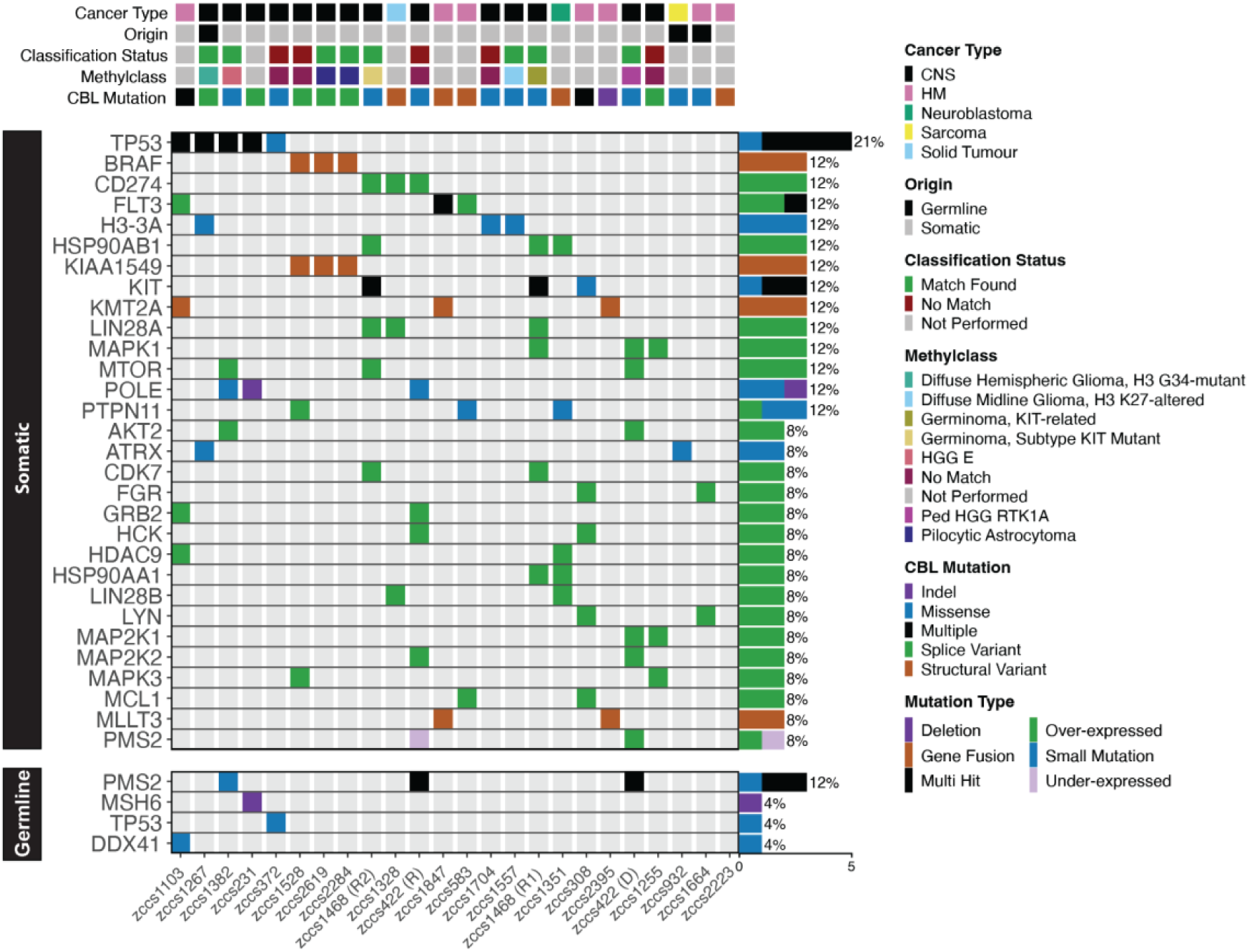
*CBL* mutations are present in molecularly diverse tumors with features of receptor tyrosine kinase activation. Oncoplot of samples with CBL linker region/ RING finger domain mutations (n=24). Each column represents an individual patient sample. Cancer type, origin (germline/somatic), methylation classification status, methylation class, and CBL mutation type are depicted in the five top rows. Methylation was not performed for non-CNS tumors (n=10) or where there was insufficient material available (n=1). The top 30 most frequently altered genes in the somatic and all germline variants, as reported by ZERO, are shown below, with frequencies shown on the right and individual sample variants colored according to the key.

### CBL ex8/9Δ is an RTK-activating variant in neuroblastoma

The CBL ex8/9Δ variant is an established dominant-negative variant that activates FLT3 in AML(20). In our cohort, we identified this variant in 2 novel tumor contexts, neuroblastoma and malignant germ cell tumor. Molecular profiling showed that the germ cell tumor (zccs1328) was highly aneuploid, with focal amplification at 12q14.3 and 12q15 including *MDM2*, and high RNA expression of *CD274*, *DNMT3B*, *LIN28A/B* and *TNFRSF8*. In the neuroblastoma patient sample (zccs1351), pathogenic SNVs: *H3C3* K28M, *MYCN* P44L, *PTPN11* A72T; *MYCN* CNV associated with Chr17q gain (3 copies); and elevated RNA expression of *GAB2*, *HDAC2/9*, *HSP90AA1/B1*, *LIN28B*, and *PHOX2A/B* was reported. A heterozygous deletion in *CBL* (chr11:119278169-119278969, hg38), resulting in CBL exon 8/9 deletion, was identified by WGS, but not reported due to insufficient evidence of pathogenicity in neuroblastoma. PCR amplification of the region spanning exons 8 and 9 of *CBL* from tumor cDNA, showed that both tumor samples (zccs1351 and zccs1328) expressed CBL WT, CBL ex8Δ, and CBL ex8/9Δ isoforms (Supplementary Fig. S4). There was sufficient neuroblastoma sample to generate a patient-derived xenograft (PDX) model and subsequently, perform high-throughput drug screening (HTS) on PDX cells. Western blot analysis confirmed expression of the CBL ex8/9Δ and WT CBL isoforms in zccs1351 PDX cells (Fig. 3A). Interestingly, this sample responded to the multi-TKI pazopanib (VEGFR, PDGFR, VGFR, KIT inhibitor) in HTS without any other associated molecular features to explain the response (Fig. 3B). Notably, this was the only neuroblastoma sample screened as part of ZERO HTS (n=15) for which pazopanib was a reported hit, suggesting that CBL mutation may be a marker of response.

**Figure 3.**
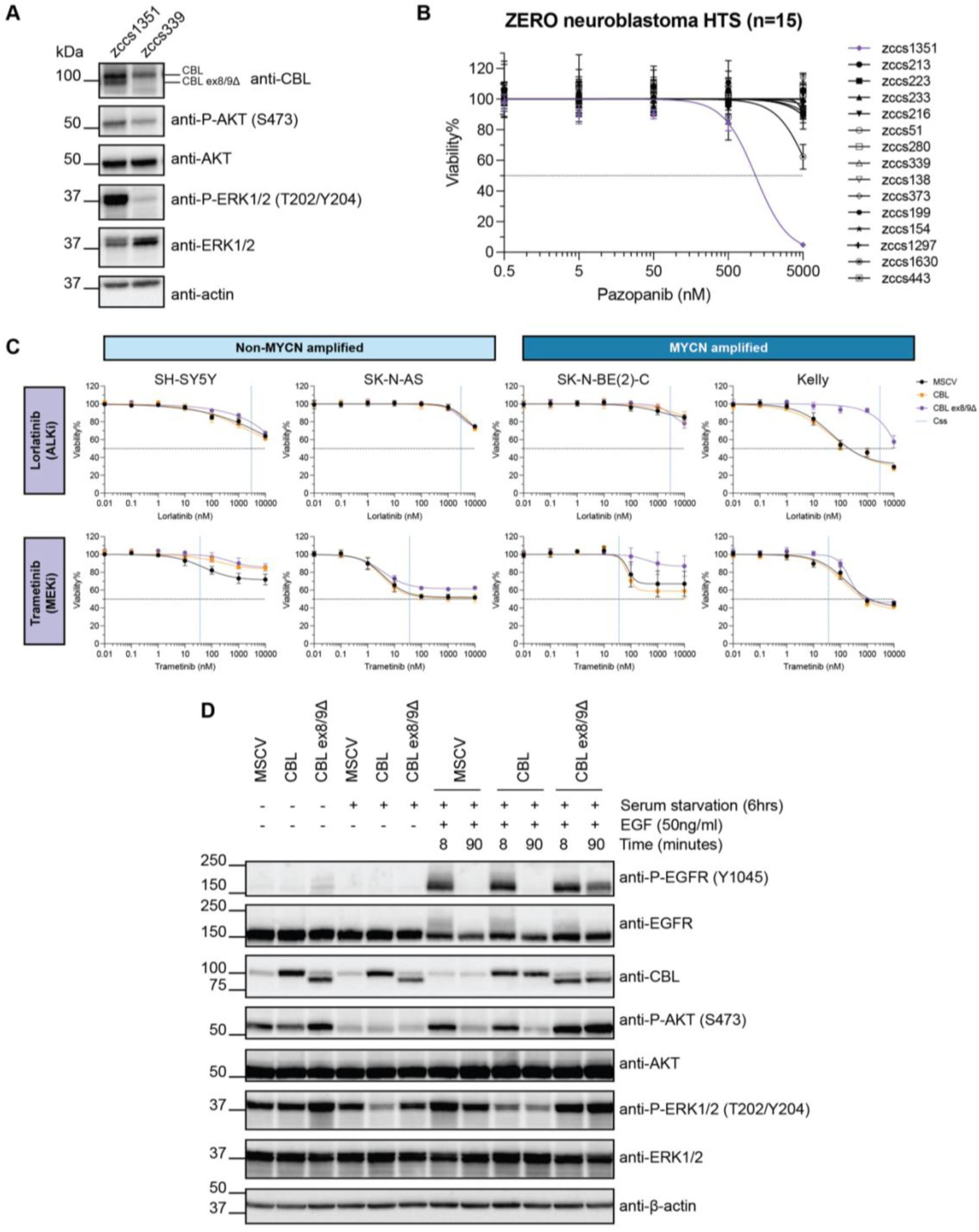
CBL exon 8/9 deletion (ex8/9Δ) is an RTK-activating variant in neuroblastoma. A) Western blot analysis of PDX cells from the CBL ex8/9Δ patient (zccs1351) or a Wt CBL control (zccs339). B) Dose response curves of neuroblastoma primary or PDX cells, screened as part of ZERO HTS, treated with pazopanib (n=15). zccs1351 is depicted by the purple line. C) Dose response curves for non-MYCN amplified (SH-SY5Y and SK-N-AS) and MYCN amplified (SK-N-BE(2)-C and Kelly) cell lines transduced with CBL variants and treated with lorlatinib (ALKi) or trametinib (MEKi). Steady-state concentration (C_ss_) of each of the drugs is represented by the blue dotted line. Starting viability for each cell line is depicted on the y-axis by the dotted lines. Cells were screened in technical triplicates in three independent experiments and data is presented as mean ± SD (n=3). D) Proliferation analysis of SH-SY5Y cell lines transduced with CBL variants. Relative viability was measured 72 hours after cell seeding and samples were normalized to MSCV control. Experiment was performed in two biologically independent cell lines in three individual experiments. Data presented as Mean ± SEM. E) Western blot analysis of EGFR signaling activation in SH-SY5Y cells stimulated with EGF for 8 or 90 minutes.

To explore the functional relevance of the CBL ex8/9Δ variant in non-hematological tumors, we cloned this variant from tumor cDNA and ectopically overexpressed CBL ex8/9Δ and WT CBL in a range of neuroblastoma cell lines. These included both non-*MYCN*-amplified (SH-SY5Y and SK-N-AS) and *MYCN*-amplified (SK-N-BE(2)-C and Kelly). The expression of CBL ex8/9Δ amplified intracellular signaling in some of these cell lines (Supplementary Fig. S5). CBL variant neuroblastoma cell lines were treated with a range of multi-(e.g. pazopanib) or specific (e.g. lorlatinib [ALK, ROS1 inhibitor] and gefitinib [EGFR inhibitor]) RTK inhibitors, as well as inhibitors targeting downstream intracellular signaling pathways (e.g. trametinib [MEK inhibitor] and paxalisib [PI3K, mTOR inhibitor]) (Supplementary Fig. S6-7). Expression of CBL ex8/9Δ rendered Kelly cells, which harbor an ALK Phe1174Leu mutation, resistant to the ALK inhibitor lorlatinib (Fig. 3C), when compared to both wild-type (WT) CBL and MSCV control cells (see S5 for IC50 values). A similar pattern was also observed with earlier generation ALK inhibitors, crizotinib and alectinib (Supplementary Fig. S7B). In addition, a varying increase in resistance to the MEK inhibitor, trametinib, was observed across all cell lines, with the most pronounced effect observed in SK-N-BE(2)-C cells (IC50 = 410nM, compared to 78nM (MSCV) and 74nM (CBL); Fig. 3C). These data suggest that CBL ex8/9Δ activated alternative signaling pathways that may drive resistance to ALK or MEK inhibitors.

Given the MYCN status of the tumor, we selected the non-MYCN amplified SH-SY5Y cell lines to further look at the impacts of CBL ex8/9Δ on CBL RING finger function and degradation of activated RTKs. To do this, we stimulated SH-SY5Y cells with EGF to activate EGFR, which is endogenously expressed in these cells. We showed that while phosphorylated EGFR was degraded by WT CBL, CBL ex8/9Δ was sufficient to block CBL mediated degradation, resulting in prolonged phosphorylation of EGFR and activation of downstream signaling, indicated by phosphorylation of AKT and ERK1/2 (Fig. 3D). These data demonstrate that in neuroblastoma cells, CBL ex8/9Δ can function to maintain activation of RTKs and represents a potential biomarker for TKI therapy inclusion in neuroblastoma.

### Alternative *CBL* splicing occurs as a result of novel *CBL* splice mutations and in the absence of genomic mutations in pediatric cancers

We identified 8 individual splice site or splice region variants in 7 patients, including 6 CNS tumors and 1 AML. Mutation of the *CBL* exon 8 splice acceptor site (c. 1096-1G) is associated with JMML(24, 26) and AML(19) and considered pathogenic, typically resulting in exon 8 skipping. We identified CBL c.1096-1G>T mutations in 3 samples: 2 pilocytic astrocytoma (zccs2284 and zccs1528) and 1 HGG (zccs231). Mutations at this site have been previously shown to result in the production of multiple *CBL* splice isoforms(26), which is consistent with observed expression of a novel dominant E366_E373del (24nt deletion) and ex8Δ isoform in RNAseq data in zccs231 (Fig. 4A-B). A likely pathogenic mutation in the exon 9 splice acceptor site (c. 1228-1G>A) was identified in a patient with radiation-induced HGG (zccs1255) and exon 9 skipping was confirmed by RNAseq. In addition, we identified 4 novel splice region variants in 3 individual patients, including 1 HGG (zccs1267), 1 pilocytic astrocytoma (zccs2619) and 1 AML (zccs308) sample, 2 of which were shown to result in exon 8 skipping in RNAseq data. In total, RNAseq data could validate the expression of alternate *CBL* isoforms in 4/6 samples, for which RNAseq was performed. To validate alternative splicing of *CBL* in zccs1528 and zccs1267, we performed PCR with forward primers spanning either the exon 7/8 or exon 7/9 boundary, to amplify WT CBL or CBL ex8Δ isoforms, respectively (Fig. 4C), which demonstrated that the CBL ex8Δ isoform is expressed at some level in these tumors. It should, however, be noted that this experiment is not quantitative, and CBL ex8Δ expression is below the limit of detection of our RNAseq, suggesting that the alternate isoform is expressed at much lower abundance compared to the WT. RNA was not available for zccs2284, so the impact of c.1096-1G>T mutation on splicing of *CBL* in this sample could not be validated by RNAseq or PCR.

**Figure 4.**
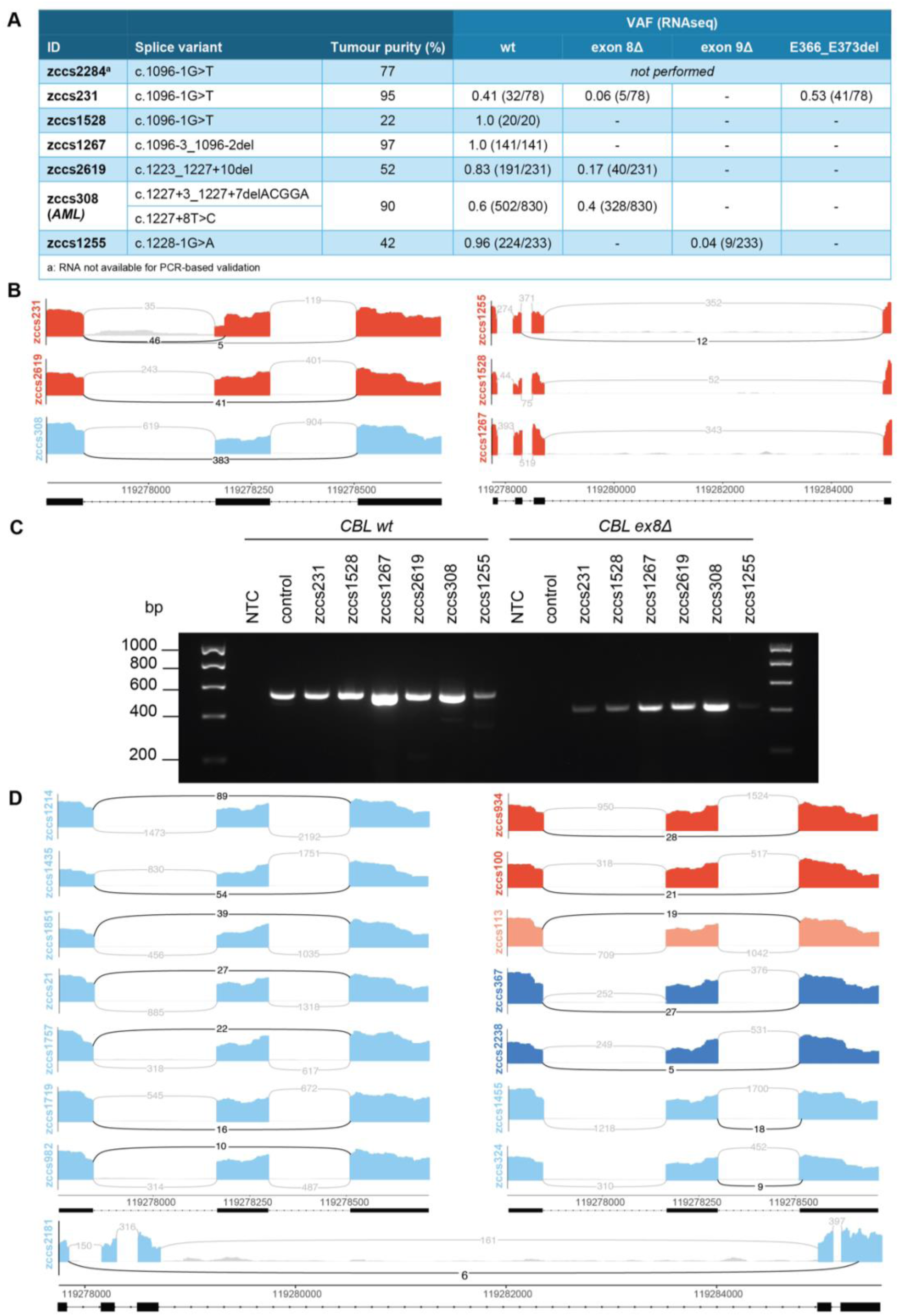
Alternative *CBL* splicing occurs as a result of novel *CBL* splice mutations and in the absence of genomic mutations in pediatric cancers. A) Table of *CBL* splice mutations. Variant Allele Frequency (VAF) for each CBL isoform was determined using RNAseq data and the number of reads for each isoform relative to the total is given in brackets. RNAseq data was not available for zccs1255. B) Sashimi plots of *CBL* alternative splicing in RNAseq data from patients with genomic *CBL* splice mutations. Exons 7-9 are depicted on the left and exons 7-10 on the right. Plots are colored according to cancer type (red = CNS, light blue = hematological malignancy). Genomic coordinates are in hg38. C) PCR analysis of CBL isoform expression. CBL wt or exon 8Δ isoforms were amplified from patient cDNA using forward primers that spanned either the exon 7/8 boundary (wt) or exon 7/9 boundary (exon 8Δ) and a reverse primer targeting exon 11, producing either 554bp or 422bp fragments, respectively. D) Sashimi plots of CBL alternative splicing in RNAseq in the absence of a splice site mutation. Exons 7-9 are depicted except in the bottom plot (zccs2181) which depicts exons 7-11. Plots are colored according to cancer type (light blue = hematological malignancy, red = CNS, peach = non-sarcomatous extracranial solid tumor, dark blue = sarcoma). Genomic coordinates are in hg38.

Next, we wanted to assess whether alternative splicing of *CBL* occurred in the absence of canonical splice site mutations. To do this we analyzed RNAseq data from ZERO for the presence of aberrant splice junctions, using an in-house method called *JuncSeek*(32), focusing on splice junctions involving exons 7-10 as the key functional regions of CBL (see methods). We identified aberrant splicing of *CBL* in 15 samples without *CBL* mutations. Most of these samples (12/15) harbored *CBL* exon 8 deletion isoforms, while the remainder had aberrant splicing in either exon 9 (n=2) or exon 11 (n=1), resulting in the partial loss of these exons (Fig. 4D). Focusing on samples with *CBL* exon 8 deletion, this occurred in 7 hematological, 2 CNS and 2 sarcoma samples (see Supplementary Table S6).

### Novel CBL variants identified in pediatric brain tumors can cooperate with RTK overexpression to drive cellular transformation

To explore the impact of novel CBL variants on cellular transformation and CBL RING finger function, we overexpressed select variants in Ba/F3 cells (Ba/F3 WT). Ba/F3 are an IL-3 dependent cell line that have been widely used to model CBL variants(20, 26, 33). We overexpressed two novel missense mutations, C384G and F378I, and the novel splice isoform, E366_E373del, all of which were identified in HGG patients, and assessed the capacity of these variants to drive IL-3 independent survival and proliferation. We used WT CBL and CBL ex8/9Δ as negative and positive controls, respectively. In short term assays, like the CBL ex8/9Δ, CBL E366_E373del was able to block IL-3 withdrawal induced cell death (Fig. 5A). Longer term assays showed that CBL E366_E373del could promote IL-3 independent cell proliferation but was slower to transform when compared to CBL ex8/9Δ (Fig. 5B). A small proportion of CBL C384G cells remained alive following IL-3 withdrawal in some experimental repeats (Fig. 5A), but these cells were not actively proliferating in longer-term assays (Fig. 5b). CBL F378I had no effect on transforming wild-type Ba/F3 cells.

**Figure 5.**
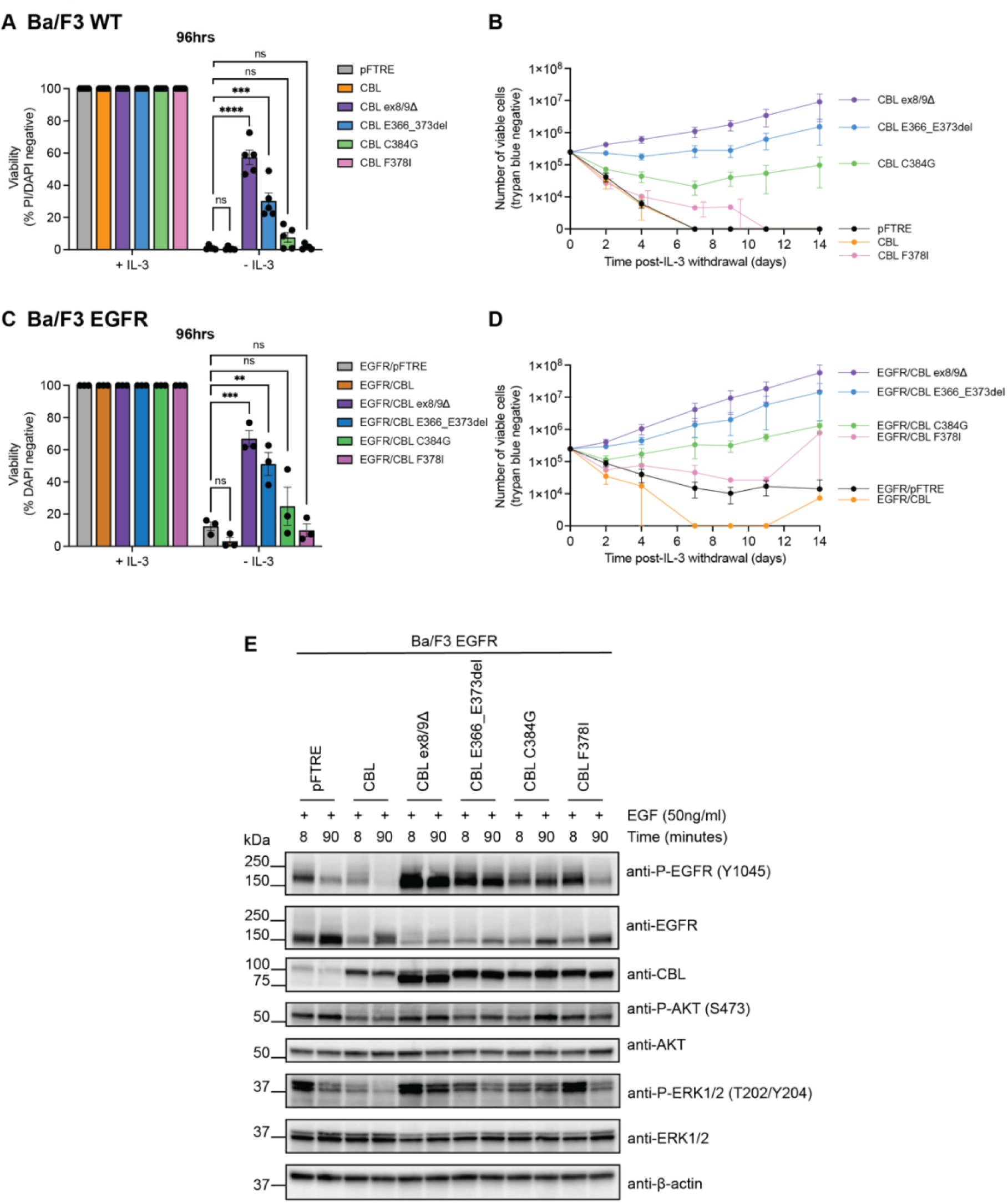
Novel CBL variants identified in paediatric brain tumors can cooperate with RTK overexpression to drive cellular transformation. A) Viability analysis of Ba/F3 cells transduced with CBL variants or an empty vector control (pFTRE), cultured in the presence (+IL-3) or absence (-IL-3) of IL-3 for 96 hours. Viability was determined by PI or DAPI exclusion, measured by flow cytometry. Data is presented as mean ± SEM (n=5). CBL variant cell lines were compared to pFTRE control using unpaired t tests with Bonferroni-Dunn correction for multiple comparisons (ns = not significant, *** = P value ≤0.001, **** = P value ≤0.0001). B) Number of viable (determined by trypan blue exclusion) Ba/F3 cells cultured without IL-3 over a 14-day period. Data is presented as mean ± SEM (n=3). C) Viability analysis of Ba/F3 cells transduced with EGFR and CBL variants, cultured in the presence or absence of IL-3 for 96 hours. Viability was determined by DAPI exclusion, measured by flow cytometry. D) Number of viable Ba/F3 EGFR cells cultured without IL-3 over a 14-day period. E) Western blot analysis of EGFR signaling activation in Ba/F3 cells, with co-expression of EGFR and CBL, stimulated with EGF for 8 or 90 minutes.

Previous studies have suggested that the loss of WT *Cbl* may be required for transformation of Ba/F3 cells by *CBL* missense variants(26, 33). The loss of WT *Cbl* is consistent with what is observed in the context of myeloid neoplasms and malignancies, where acquired uniparental disomy and loss of chromosome 11q results in the loss of the normal *CBL* allele(22–25). In contrast, variants that delete all or part of CBL exon 8, which contains the critical linker region, do not require WT CBL loss and can transform Ba/F3 cells when co-expressed with an RTK (for example, FLT3(20)). In our cohort, CBL variants were not always associated with WT *CBL* loss, and were more frequently heterozygous or sub clonal (Supplementary Table S1). Thus, we assessed the capacity of our variants to cooperate with overexpression of an RTK in the presence of WT CBL. To do this, we generated a Ba/F3 cell line with doxycycline-inducible overexpression of EGFR (Ba/F3 EGFR) and transduced these cells with each CBL variant. These cells were then subjected to the same IL-3 withdrawal assay performed on Ba/F3 WT. As was observed in Ba/F3 WT, CBL E366_E373del and CBL ex8/9Δ were able to block short term IL-3 withdrawal induced cell death in Ba/F3 EGFR cells (Fig. 5C). Notably, EGFR overexpression alone (pFTRE control) was sufficient to promote survival in a small proportion of cells in the absence of IL-3, which was similar to what was observed in cells expressing the C384G and F378I missense mutations. In longer term assays, CBL E366_E373del, CBL ex8/9Δ and CBL C384G consistently transformed Ba/F3 EGFR cells and promoted proliferation, while cells expressing F378I or EGFR alone (pFTRE) stopped proliferating despite early signs of transformation (Fig. 5D).

At baseline, overexpression of EGFR and CBL E366_E373del, mediated by doxycycline addition, consistently promoted maintained activation of EGFR signaling (marked by phosphorylation), like CBL ex8/9Δ (Supplementary Fig. S8). This was not observed for the CBL F378I mutant, while the impact of CBL C384G was variable. To specifically look at the impact of CBL variants on degradation of activated EGFR, we stimulated Ba/F3 EGFR cells with EGF ligand and analyzed EGFR phosphorylation and downstream signaling pathways (Fig. 5E). Overexpression of CBL ex8/9Δ, CBL E366_E373del, and to some extent CBL C384G and CBL F378I, blocked degradation of phosphorylated EGFR, as observed in cells expressing WT CBL. Sustained activation of downstream signaling, marked by expression of phosphorylated AKT and ERK protein expression, was observed with all variants, albeit to varying degrees, in comparison to WT CBL. Taken together, these data demonstrate that novel CBL missense mutants and splice isoforms identified in pediatric brain tumors can cooperate with RTK overexpression to drive transformation, suggesting that these variants may represent RTK-activating and therapeutically targetable events.

## Discussion

Molecularly targeted therapies and precision medicine have provided therapeutic options for pediatric patients with high-risk and difficult-to-treat cancers, improving individual outcomes(4). Further improvements depend on more effective targeting of existing druggable or currently undruggable targets, or the identification of novel cancer driver genes that may enable more patients to be treated with existing therapies. Here, we have identified *CBL* mutations in novel tumor contexts: pediatric solid and CNS tumors. *CBL* mutations include novel missense and splice mutations, in addition to established *CBL* driver variants. *CBL*-mutated tumors are molecularly diverse, but have features of RTK activation, supporting the proposition that CBL mutations may cooperate with either mutated or wild-type RTKs to enhance RTK activation and signal transduction. We provide functional evidence that established *CBL* variants are RTK activating in novel tumor contexts and that novel *CBL* variants can cooperate with RTKs to drive transformation in cytokine dependent models. This work is the first to comprehensively characterize *CBL* variants in non-hematological malignancies and suggests that *CBL* variants may represent a biomarker for TKI therapy in other tumor contexts.

The genomic analysis of pediatric tumors performed as part of large pan-cancer genomic studies(34–36), or in the context of precision medicine(4, 37–39), have enabled greater understanding of the molecular drivers of childhood cancers. *CBL* mutations are well-established in the context of myeloid malignancies, including myeloproliferative neoplasms and AML(19, 20, 23), and have also been identified in mixed-phonotype and lymphoid leukemias(34, 36). In the pediatric setting, germline *CBL* mutations characterize CBL syndrome, a Noonan syndrome-like RASopathy, and predispose patients to JMML(26). *CBL* mutations are rarely described in other tumor types in international pediatric cancer cohorts. The Genome for Kids study identified a germline *CBL* splice site mutation in a patient with a rare type of malignant germ cell tumor, dysgerminoma, with an FGFR2 R253R missense mutation(36). This *CBL* splice mutation was homozygous in the tumor due to copy neutral Loss of Heterozygosity (LOH) at chr11 and resulted in exon 9 skipping. In our cohort, we identified CBL ex8/9Δ in a testicular germ cell tumor, as well as a novel CBL D388Y mutation in 2 relapsed samples from a CNS germinomatous germ cell tumor. In publicly available datasets, including St Jude Cloud(40) and PedcBioPortal(41), *CBL* mutations are found rarely in non-hematological tumors and include missense, indels and splice mutations (a total of 16 patient samples). Missense mutations have been identified in single cases of HGG or low-grade glioma (LGG) including Y368C, Y371H, C381R, C381Y, and E386K, as well as an additional case harboring two variants (Y368C and C401F). One deletion, L405del, was identified in a HGG sample. *CBL* splicing variants are however, more frequently identified, including the established pathogenic splice acceptor variants c.1096-1G>T (n=6 samples; 4x LGG, 1x ependymoma and 1 x osteosarcoma) and c.1096-2A>T (n=1, ependymoma). Notably, *CBL* mutations frequently co-occur with the *KIAA1549-BRAF* fusion in these datasets (n=5), consistent with our findings of 3 additional cases. While rare, *CBL* mutations have been identified in other pediatric pan-cancer cohorts, but the relevance of these mutations in non-hematological tumors had remained unappreciated as a result of the unusual tumor contexts. It is also likely that *CBL* mutations occur at an even higher frequency but are being filtered out in other datasets using predefined gene lists for each cancer type.

Our study indicates that *CBL* mutations are most frequently associated with CNS tumors, particularly in pHGG and LGG. The comprehensive molecular approach in ZERO (WGS and RNAseq) enabled the identification of splice mutations, both canonical splice acceptor/donor mutations and novel splice region mutations, that could be missed by other methodologies such as whole exome or panel sequencing(42). We, and others(26), have demonstrated that *CBL* splice mutations result in the production of a range of aberrant *CBL* transcripts lacking exon 8, exon 9 or parts of either exon. In addition to the validation of splicing outcomes resulting from genomic variants, we showed that while rare, aberrant *CBL* transcripts lacking exon 8 also arise in the absence of *CBL* splice region variants. Since deletion of *CBL* exon 8 is pathogenic and CBL exon 8Δ is functionally transforming(20), the expression of this transcript, even in the absence of known explanatory mutations may be a contributory and targetable driver in such cases. This further highlights the value of combined whole genome and transcriptomic sequencing to capture splicing variants and aberrantly spliced transcripts.

There are conflicting perspectives on the requirement for loss of wild-type *CBL* for *CBL* mutations to drive cancer. The association of *CBL* mutations with loss of wild-type CBL in the context of 11q loss of uniparental disomy in myeloid malignancies has reinforced this notion(21–23). A paradigm was established based on early functional studies suggesting that there are two classes of *CBL* mutation(16). The first being transforming variants, which include single amino acid or larger intragenic deletions impacting the alpha-helix of the linker region of CBL (V363_G375) and are thought to act in a dominant-negative manner. The second being missense mutations or deletions that impact the RING finger domain, that while sufficient to abolish CBL-mediated polyubiquitination, cannot drive cell transformation. The caveat to this model is the potential role of cooperating molecular events. Our data provide support for the notion that deletion of wild-type Cbl is not required for overexpressed mutant Cbl to drive a phenotype of cytokine-independent proliferation and repression of apoptosis. We have shown that while deletion variants (CBL exon 8/9Δ and CBL E366_E373del) could drive cell transformation in wild-type Ba/F3 cells, CBL missense mutations were insufficient, supporting previous reports that wild-type *Cbl* loss is required in this model. However, when co-expressed with EGFR, CBL C384G was able to drive cell transformation, suggesting that CBL missense mutations can cooperate with RTK overexpression to drive transformation(18, 22).

The potential therapeutic implications of *CBL* mutation are currently dependent on an understanding of which RTK is dysregulated. FLT3 inhibitors have demonstrated some efficacy in CBL-mutant leukemia cell lines *in vitro* and *in vivo*(43). FLT3 inhibitors — midostaurin, gilteritinib, and quizartinib — are currently used clinically for the treatment of AML, but their efficacy against CBL mutation has not been established, as clinical trials have primarily focused on FLT3 mutation status(44). A Phase I clinical trial assessing quizartinib in combination with azacitidine in treatment naive or relapsed/refractory MDS/MPN with either *FLT3* or *CBL* mutations showed some response in CBL-mutated patients (60% overall response rate (ORR); n=5), although this was not as high as FLT3-ITD patients (100% ORR; n=7) The Phase II trial is currently ongoing and will enable greater understanding of the potential efficacy of this combination in CBL-mutated AML. The target RTK in the context of pediatric brain tumors remains unknown. As in AML, it is likely that mutant CBL functions to enhance activity of wild-type RTKs. Mutant CBL has been shown to have no effect on FLT3-ITD regulation(45), and most commonly occurs in the context of wild-type *FLT3*(20, 22). The majority of CBL mutant samples in our cohort did not express genomic mutations in RTK genes, supporting the hypothesis that mutant CBL would function to amplify signaling from wild-type RTKs in these tumors. However, the co-occurrence of *CBL* splicing mutations with the *KIAA1549::BRAF* fusion in LGG in our cohort and others raises the possibility that CBL may function to amplify KIAA1549::BRAF signaling in these cases. To identify the target RTK in individual tumors, (phospho)proteomic approaches are likely to be the most valuable. Future work will focus on modelling CBL mutations in HGG patient-derived cell lines and neural stem cell models to elucidate potential target RTKs in this cancer context. It should be noted that Cbl-b inhibitors have recently been developed as an immunotherapy agent in cancer, and function to enhance T-cell activation and promote antitumor activity(46). The ability of these inhibitors to target mutant c-Cbl and therefore the therapeutic benefit in CBL-mutated cancers remains unknown.

In all, this study demonstrates that *CBL* mutations, both known and novel, occur in pediatric solid and CNS tumors. We show that unbiased comprehensive molecular profiling approaches can identify novel cancer drivers and is particularly valuable for the identification and validation of splice mutations. Our findings highlight broader implications for precision medicine approaches and reiterate the need to consider molecular alterations identified in atypical cancer contexts, as these may be functional and actionable. Our early functional data demonstrates that established CBL variants are functional in non-hematological tumors and characterizes novel CBL variants in established cytokine-dependent model systems. To realize the potential therapeutic benefit of this finding, future work will focus on the identification of the target RTK in the context of specific cancer types or individual patient tumors. This will reveal novel drug targets for pediatric cancer patients, enabling more patients with aggressive and difficult-to-treat cancers to receive targeted therapies.

## Methods

### Molecular profiling

Molecular profiling (whole genome sequencing, RNA sequencing and DNA methylation analysis) was obtained from the ZERO program, as previously described(4). Data obtained was aligned to the human genome reference hg38 and annotated with gencode v41. Additional RNA sequencing and methylation analysis were performed as described below. Somatic and germline variants of CBL mutated samples were plotted using ggoncoplot(47).

### Clustering Analysis

For both RNA sequencing and methylation, unsupervised clustering analysis was performed in R using t-SNE (t-Distributed Stochastic Neighbour Embedding) analysis with the Rtsne (v0.17; https://github.co/jkrijthe/Rtsne) package. Cohort wide analysis was only conducted on RNA sequencing data (N=1653). Both RNA sequencing and methylation data was filtered to only samples with a CNS diagnosis, resulting in CNS specific clustering on N=589 RNA sequencing samples and N=627 methylation samples.

### RNA sequencing splice isoform analysis

For the analysis of aberrant *CBL* splice isoforms, we used JuncSeek(32), developed in-house for the identification of aberrant splice junctions in RNAseq data. We focused on the region of *CBL* encoding the linker and RING finger regions (exon 7-10) and filtered the data based on start_exon7>X and end_exon10<X to extract only splice junctions that occurred in the region of interest (hg38 chr11:119,277,756-119,285,099) and ≥5 reads of evidence supporting the junction. The presence and structure of splice isoforms were then validated through manual inspection of RNAseq reads in Integrative Genomics Viewer (v. 2.16.2, Broad Institute).

### Protein modelling and pathogenicity prediction

Protein variants pathogenicity was modelled using the ESM family of protein language models. Briefly, the amino acid sequence of human CBL (canonical isoform) was used as input to ESM-2, and for each position we obtained the log-likelihood for all 20 possible residues. For visualization of predicted positional constraint, we calculated a residue-wise log-likelihood ratio (LLR) as the difference between the log-likelihood of each possible amino acid and that of the wild-type residue at that position and plotted the resulting LLR matrix across the length of the protein. To visualize the structural context of these variants, we generated a three-dimensional model of CBL using ESMfold, and mapped each mutant residue onto the model, with variant color intensity indicating the pathogenicity score. To estimate the effect of individual CBL missense variants, we extracted from the LLR matrix the score corresponding to the mutant residue at the relevant position and used this as the pathogenicity score; variants with absolute scores ≥ 7.5 were classified as “pathogenic.”

### Ethics and patient samples

This study was approved by the University of New South Wales Human Research Ethics Committee (Number: iRECS7552). Access to patient samples was approved and obtained through the ZERO Childhood Cancer Program Research Management Committee.

### *CBL* variant cloning and DNA constructs

*CBL* variant constructs were either cloned from patient material obtained from the ZERO program. Patient RNA was first reverse transcribed into cDNA using the Invitrogen™ SuperScript™ III First-Strand Synthesis System (Thermo Fisher Scientific, #18080051). Full length *CBL* variant cDNA fragments were amplified from patient cDNA using Q5^®^ Hot Start High-Fidelity DNA Polymerase (New England Biolabs, #M0493) and the following primers targeting the 5’ and 3’ ends of *CBL*: BamHI CBL Forward: 5’ AATGGATCCATGGCCGGCAACGTGAAG and NheI CBL Reverse: 5’ AATGCTAGCCTAGGTAGCTACATGGGCAGG. As a control, primers targeting exon 6 and 11 were designed to amplify the variant/deleted region (exon 8-9) of CBL, as follows: CBL del Forward: 5’ CTGAGCTGTACTCGTCTGGG and CBL del Reverse: 5’ CTAGGTAGCTACATGGGCAGG. All variants (CBL WT, ex8/9Δ, E366_E373del, C384G and F378I) were cloned by restriction digest into the pFTRE GFP lentiviral vector using BamHI and NheI restriction sites according to standard procedures. CBL WT and CBL ex8/9Δ cDNA fragments were subsequently PCR amplified from plasmid DNA using the following primers: BamHI CBL Forward (sequence above) and MluI CBL Reverse: 5’ AATACGCGTCTAGGTAGCTACATGGGCAGG, and sub-cloned into the MSCV-IRES-GFP vector (MSCV) using BamHI and MluI restriction sites.

For the generation of the EGFR overexpression plasmid, sub-cloned into the pFTRE tdTomato lentiviral vector(5) from the EGFR WT pBABE Puro plasmid (Addgene, #11011). EGFR was PCR amplified from plasmid DNA using the following primers: NheI EGFR Forward: 5’ ATAATGCTAGCATGCGACCCTCCGGGACG and NheI EGFR Reverse: 5’ TTCCAGCTAGCTCATGCTCCAATAAATTCAC and sub-cloned into pFTRE tdTomato using the NheI using standard procedures.

DNA construct sequences were confirmed by Sanger sequencing (Ramaciotti Centre for Functional Genomics, UNSW).

### PCR analysis of CBL variant splicing outcomes

To validate the impact of genomic *CBL* splicing variants on *CBL* splicing, we performed PCR to specifically amplify either WT *CBL* or *CBL* ex8Δ transcripts. PCR was performed on cDNA from 7/8 patients (RNA was not available for one patient, zccs2284) with genomic *CBL* splice mutations Q5^®^ Hot Start High-Fidelity DNA Polymerase. For amplification of the WT *CBL* isoform the following primers were used: CBL ex7/8del Forward: 5’ AAAGTGACCCAGGAACAATATGAA and CBL del Reverse: 5’ CTAGGTAGCTACATGGGCAGG (and as per “*CBL* variant cloning and DNA constructs”). For amplification of the *CBL* ex8Δ isoform the following primers were used: CBL ex7/9 Forward: 5’ AAAGTGACCCAGGAATCAGAAGGT and CBL del Reverse (as above). cDNA from a patient with WT *CBL* was used as a negative control.

### Cell culture and retroviral and lentiviral transduction

HEK293T, SH-SY5Y, SK-N-AS and SK-N-BE(2)-C cells were maintained in DMEM (Thermo Fisher Scientific, #11995-065) supplemented with 10% fetal bovine serum (FBS, Thermo Fisher Scientific, #10100147), at 37°C and 5% CO_2_. Ba/F3 cell lines were maintained in RPMI 1640 medium (Thermo Fisher Scientific, #22400-089) supplemented with 10% FBS and 0.5 ng/mL mouse interleukin-3 (IL-3) recombinant protein (Thermo Fisher Scientific, #213-13-10), at 37°C and 5% CO_2_.

Retroviral and lentiviral transfection of HEK293T cells and transduction of neuroblastoma cell lines (SH-SY5Y, SK-N-AS, SK-N-BE(2)-C and Kelly), and Ba/F3 and zccs1647, with CBL MSCV or CBL pFTRE GFP constructs, respectively, was performed using Effectene Transfection Reagent (QIAGEN, #301425), as previously described(48). Transduced cell lines were sorted for GFP or GFP/dTomato expression by fluorescence-activated cell sorting (FACS) using the FACSAria™ Fusion (BD Biosciences).

### EGF stimulation

For SH-SY5Y cells, cells were counted and seeded at 7.4 x 10^5^ viable cells in a 10cm tissue culture dish and incubated for 96 hours. Following incubation, complete growth medium was aspirated, cells were washed with PBS, replenished with 10ml of serum-free DMEM (except for untreated control), and incubated for 6 hours prior to EGF stimulation. Cells were subsequently stimulated with 50ng/ml of EGF for either 8 or 90 minutes, or vehicle control. Following stimulation, media containing EGF was removed, and cells were washed twice in cold PBS. To dislodge cells from plates, 2ml of cold PBS was added to plates and cells were scraped off into a 15ml centrifuge tube using a cell scraper (Grenier, #541070). Cell pellets were harvested according to standard procedures.

For Ba/F3 EGFR cells, CBL variant expression was induced with doxycycline 72 hours prior to growth factor withdrawal and stimulation (day 0). Following 72-hour incubation, cells were washed three times in PBS and resuspended in mIL-3-free medium for counting, as described above. Cells were seeded at 1.5 x 10^6^ cells in 2ml of media in a 12-well plate for a final concentration of 7.5 x 10^5^ cells/ml and incubated for 6 hours in IL-3-free conditions. Cell pellets from EGF stimulation assays were lysed and quantified, as described below, prior to Western blotting.

### Western blotting

Whole cell lysates were isolated by lysing cells in RIPA Lysis and Extraction Buffer (Thermo Fisher Scientific, #89901) according to standard procedures. Protein was quantified using the Pierce™ BCA Protein Assay (Thermo Fisher Scientific, #23225), according to the Manufacturer’s instructions. 30-50ug of protein was separated by SDS-PAGE under denaturing conditions, and proteins were transferred onto Amersham^™^ Protran^®^ nitrocellulose membrane (Merck, #GE10600016) using the Criterion™ or Mini-PROTEAN Tetra Cell systems according to the Manufacturer’s instructions (Bio-Rad Laboratories). Western blotting was performed with primary and secondary antibodies detailed in Table 2. Antibodies were detected using Clarity™ (Bio-Rad Laboratories, #1705061) or Clarity Max™ (Bio-Rad Laboratories, #1705062) Western ECL Substrates and the Chemidoc™ Imaging System (Bio-Rad Laboratories). Chemiluminescent images were analyzed using Image Lab (v6.1.0, Bio-Rad Laboratories).

**Table 2.** Western blot antibodies.

| **Antibody** | **Supplier** | **Catalogue number** | **RRID** |
| --- | --- | --- | --- |
| Phospho-ALK (Tyr1278) | Cell Signaling Technology | 6941 | AB_10860598 |
| ALK | Cell Signaling Technology | 3633 | AB_11127207 |
| EGFR (phospho Tyr1045) | Cell Signaling Technology | 2237 | AB_331710 |
| EGFR | Cell Signaling Technology | 4267 | AB_2246311 |
| c-CBL | Cell Signaling Technology | 2747 | AB_2275284 |
| Phospho-AKT (Ser473) | Cell Signaling Technology | 4060 | AB_2315049 |
| AKT | Cell Signaling Technology | 4691 | AB_915783 |
| Phospho-ERK1/2 (Thr202/Tyr204) | Cell Signaling Technology | 9101 | AB_331646 |
| ERK1/2 | Cell Signaling Technology | 4695 | AB_390779 |
| β-actin | Sigma Aldrich | A2228 | AB_476697 |
| Anti-rabbit HRP-conjugated | Cytiva | NA9340 | AB_772191 |
| Anti-mouse HRP-conjugated | Cytiva | NA931 | AB_772210 |

### Drug screening

High-throughput drug screening on the CBL ex8/9Δ neuroblastoma PDX (zccs1351) and other primary patient samples and PDXs was performed as part of the ZERO program(49).

Drug screening of neuroblastoma cell lines and data analysis were performed as previously described(5). In brief, neuroblastoma cell lines expressing WT CBL, CBL ex8/9Δ or empty vector control (MSCV), were seeded at 2-5 x 10^3^ cells per well in a 384 well plates, using the Multidrop Combi Reagent Dispenser (Thermo Fisher Scientific). Cells were incubated for 24 hours prior to drug addition, using the HP D300 Digital Dispenser (Tecan) and then treated with a 7-point dose titration 0.01-10,000nM (10-fold serial dilutions) of 12 selected kinase inhibitor drugs (MedChemExpress; see Table 3) for 72 hours. Cell viability was measured using the CellTiter-Glo® 2.0 Cell Viability Assay (Promega, #G9242) and luminescence values were captured using the EnVision and EnSpire Alpha plate readers (PerkinElmer). Drug testing was performed in technical triplicates in three independent repeat experiments (n=3). Cell viability values were used to generate dose response curves using nonlinear regression analysis and determine half-maximal inhibitory values (IC50). Data were analyzed using GraphPad Prism Software (version 10.2.3, GraphPad Prism Software, LLC).

**Table 3.** Kinase inhibitors.

| **Drug name** | **Target** | **Supplier** | **Catalogue number** |
| --- | --- | --- | --- |
| Pazopanib | VEGFR1/2/3; PDGFRA/B; FGFR1; c-Kit; c-FMS | MedChemExpress | HY-10208 |
| Regorafenib | VEGFR1/2/3; PDGFRB; c-KIT; RET; Raf-1 | MedChemExpress | HY-10331 |
| Lenvatinib | VEGFR1/2/3; PDGFRA/B; FGFR1 | MedChemExpress | HY-10981 |
| Dasatinib | Abl; Src; c-Kit; LCK; YES; FYN; PDGFRA/B | MedChemExpress | HY-10181 |
| Gefitinib | EGFR | MedChemExpress | HY-50895 |
| Afatinib | EGFR; HER2 | MedChemExpress | HY-10261 |
| Crizotinib | ROS1; c-MET; ALK | MedChemExpress | HY-50878 |
| Alectinib | ALK; RET | MedChemExpress | HY-13011 |
| Lorlatinib | ALK; ROS1 | MedChemExpress | HY-12215 |
| Trametinib | MEK | MedChemExpress | HY-10999 |
| Ulixertinib | ERK1/2 | MedChemExpress | HY-15816 |
| Paxalisib | PI3Ka/b/d/g; mTOR | MedChemExpress | HY-19962 |

### IL-3 withdrawal assays

IL-3 withdrawal assays in Ba/F3 cells were performed as previously described(5). In brief, Ba/F3 cells (WT or EGFR) transduced with CBL variants in pFTRE GFP were washed 3 times in PBS and resuspended in mIL-3-free RPMI + 10% FBS. 5 x 10^5^ viable cells (determined using a hemocytometer and trypan blue exclusion) were seeded in 2ml of +IL-3 (normal conditions; RPMI + 10% FBS + 0.5ng/ml mIL-3) or -IL-3 (RPMI + 10% FBS), for a final cell concentration of 2.5 x 10^5^ viable cells/ml, with 1ug/ml doxycycline (Merck, #D9891-1G) to induce expression of CBL variants. Cells were incubated for a total of 14 days and split to prevent overgrowth. Doxycycline was replenished every 3-4 days. The number of viable cells and cell viability were determined at 2, 4, 7, 9, 11, and 14 days by trypan blue exclusion and DAPI (Merck, #MBD0015) or propidium iodide (PI; Thermo Fisher Scientific, #P3566) exclusion, measured by flow cytometry, respectively. CBL variant expression was induced with doxycycline 72 hours prior to assay setup.

## Data Availability

All data generated or analyzed during this study are included in this published article and its supplementary information files. Raw data files are available upon request from the corresponding author. RNAseq and WGS data were generated as part of the ZERO program and are available upon request to https://www.zerochildhoodcancer.org.au/clinicians-researchers/for-researchers/data-and-sample-resources

## Acknowledgments

We gratefully acknowledge the Zero Childhood Cancer Program, Clinical Curation Team, Bioinformatics, and Computational Biology Group for providing molecular data and analysis for this study. We thank Dr Maria Tsoli for providing patient-derived HGG cell lines and expertise. We thank Tour de Cure, Luminesce Alliance, Audi Foundation, Yellow Diamond Foundation and the NHMRC (Ideas Grant 2028235. Recovering lost therapeutic opportunities in high-risk paediatric cancer - novel drivers of receptor tyrosine kinase activation) for providing funding. The authors acknowledge the provision of computing and data resources provided by the Australian BioCommons Leadership Share (ABLeS) program to perform the splice junction and clustering analysis of the RNAseq data. This program is co-funded by Bioplatforms Australia (enabled by NCRIS), the National Computational Infrastructure and Pawsey Supercomputing Research Centre.

