## Supplementary Table S5 and Supplementary Figure S1-S8 for "CBL mutations in pediatric solid and CNS tumors are a marker of receptor tyrosine kinase activation and a potential therapeutic target"

### Supplementary Tables

#### Supplementary Table S5. IC50 values from kinase inhibitor drug screen in neuroblastoma cell lines with CBL variants

| | MSCV | CBL | CBL ex8/9Δ |
|---|---|---|---|
| | ***Mean IC50 (nM) [95% CI]*** | | |
| | **SH-SY5Y** | | |
| **Pazopanib** | 41005 [*undefined,* 73451] | 145470 [24087, *undefined*] | 568751 [24087, *undefined*] |
| **Regorafenib** | 1837 [1533, *undefined*] | 1873 [1231, *undefined*] | 2005 [1633, *undefined*] |
| **Lenvatinib** | *Unstable* | *Unstable* | 6679 [1850, *undefined*] |
| **Dasatinib** | *Unstable* | 6997 [1258, *undefined*] | 1051 [144, *undefined*] |
| **Gefitinib** | 1809 [700, *undefined*] | 5888 [1234, *undefined*] | *Unstable* |
| **Afatinib** | 2076 [1823, *undefined*] | 1878 [1601, *undefined*] | 2691 [1928, *undefined*] |
| **Crizotinib** | 985 [865, *undefined*] | 979 [869, *undefined*] | 1226 [*undefined*] |
| **Alectinib** | 1233 [889, 2418] | 1012 [552, 4956] | 911 [590, 1987] |
| **Lorlatinib** | 17713 [553, *undefined*] | 1724 [238, *undefined*] | *Unstable* |
| **Trametinib** | 49.1 [19.9, 155] | 174 [36.5, 7743] | 480 [73.3, *undefined*] |
| **Ulixertinib** | 4109 [1210, *undefined*] | 4203 [1207, *undefined*] | 1644 [1035, *undefined*] |
| **Paxalisib** | 1062 [816, 1769] | 778 [625, 982] | 776 [496, 1527] |
| | **SK-N-AS** | | |
| **Pazopanib** | *Unstable* | 1184322 [53069, *undefined*] | 34406 [*undefined*, 78.8] |

| | | | |
|---|---|---|---|
| **Regorafenib** | 1201 [1068, *undefined*] | 1224 [1073, *undefined*] | 1168 [1017, *undefined*] |
| **Lenvatinib** | *Unstable* | *Unstable* | *Unstable* |
| **Dasatinib** | *Unstable* | *Unstable* | *Unstable* |
| **Gefitinib** | *Unstable* | *Unstable* | 26715 [*undefined*] |
| **Afatinib** | 2204 [1822, *undefined*] | 1946 [1722, *undefined*] | 2361 [1731, *undefined*] |
| **Crizotinib** | 1269 [*undefined*, 1369] | 1125 [*undefined*] | 1281 [*undefined*, 1384] |
| **Alectinib** | 1181 [*undefined*] | 1116 [*undefined*] | 1237 [*undefined*] |
| **Lorlatinib** | 3732 [1986, *undefined*] | *Unstable* | 2783 [1748, *undefined*] |
| **Trametinib** | 4.0 [2.6, 6.0] | 3.5 [2.2, 5.4] | 3.7 [2.4, 5.6] |
| **Ulixertinib** | 481 [314, 725] | 602 [353, 1322] | 395 [239, 663] |
| **Paxalisib** | 851 [670, 1158] | 825 [666, 1071] | 724 [576, 933] |
| | **SK-N-BE(2)-C** | | |
| **Pazopanib** | *Unstable* | *Unstable* | *Unstable* |
| **Regorafenib** | *Unstable* | *Unstable* | 6097 [*undefined*] |
| **Lenvatinib** | *Unstable* | *Unstable* | *Unstable* |
| **Dasatinib** | *Unstable* | *Unstable* | *Unstable* |
| **Gefitinib** | *Unstable* | *Unstable* | *Unstable* |
| **Afatinib** | *Unstable* | *Unstable* | 8274 [3143, *undefined*] |
| **Crizotinib** | 2503 [1456, *undefined*] | 2033 [1468, *undefined*] | 6371 [*undefined*] |
| **Alectinib** | 11997 [1094, *undefined*] | 5081 [1381, *undefined*] | 2830 [1094, *undefined*] |
| **Lorlatinib** | 1371 [148, *undefined*] | *Unstable* | 3798 [1709, *undefined*] |
| **Trametinib** | 78.1 [19.2, *undefined*] | 74.1 [44.1, *undefined*] | 410 [16.5, *undefined*] |
| **Ulixertinib** | *Unstable* | *Unstable* | *Unstable* |
| **Paxalisib** | 1624 [1389, *undefined*] | 1650 [1394, *undefined*] | 1567 [1267, *undefined*] |
| | **Kelly** | | |
| **Pazopanib** | 211797 [6160, *undefined*] | *Unstable* | 98880 [16990, *undefined*] |
| **Regorafenib** | 1617 [1358, *undefined*] | 1548 [1265, *undefined*] | 1587 [1259, *undefined*] |
| **Lenvatinib** | 511 [323, 777] | 456 [231, 974] | 754 [388, 1665] |
| **Dasatinib** | *Unstable* | *Unstable* | 334376 [*undefined*] |
| **Gefitinib** | 8494 [1590, *undefined*] | 19477 [2126, *undefined*] | *Unstable* |
| **Afatinib** | 1868 [1686, *undefined*] | 9517 [*undefined*] | 4019 [1845, *undefined*] |
| **Crizotinib** | 793 [669, 973] | 783 [647, 994] | 2837 [*undefined*, 76560] |
| **Alectinib** | 484 [268, 1163] | 477 [311, 850] | 1547 [1135, 2864] |

| | | | |
|---|---|---|---|
| **Lorlatinib** | 48.4 [26.0, 119] | 51.6 [24.6, 173] | *Unstable* |
| **Trametinib** | 160 [94.4, 318] | 139 [92.6, 227] | 221 [*undefined*, 327] |
| **Ulixertinib** | 790 [538, 1560] | 871 [486, 3644] | 2001 [1338, 31913] |
| **Paxalisib** | 390 [316, 477] | 357 [284, 445] | 550 [459, 650] |

### Supplementary Figures

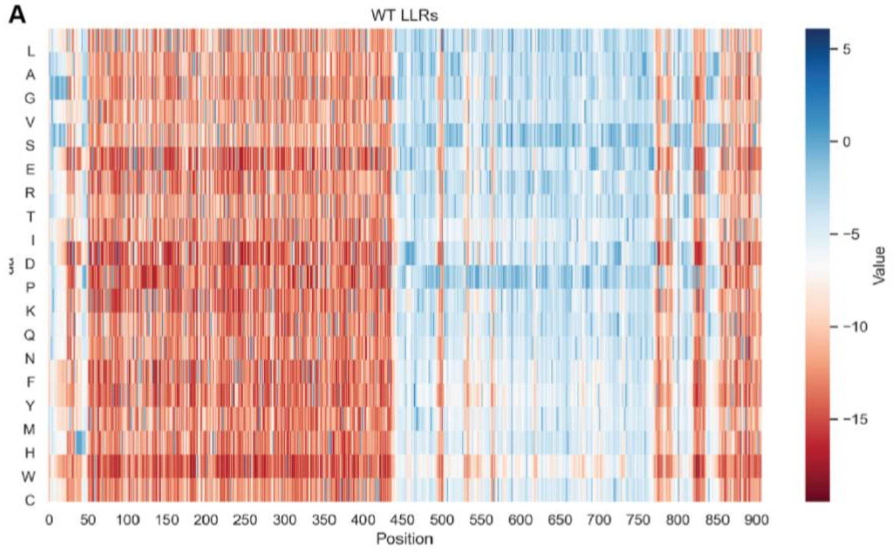


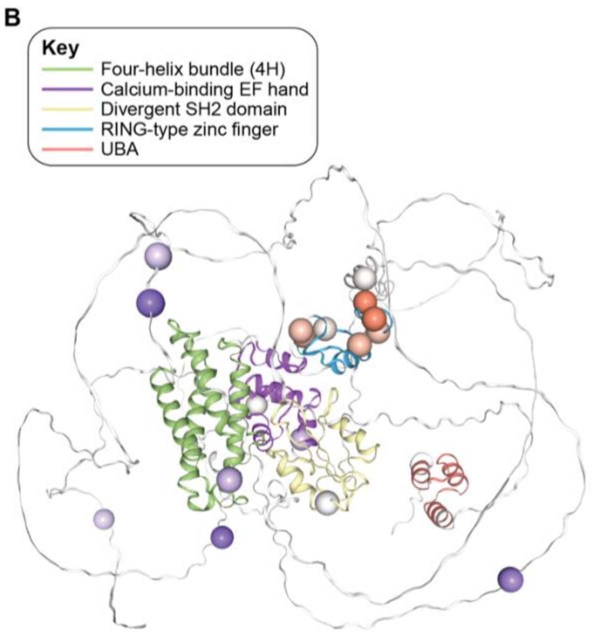


C

| Mutation | Pathogenicity score |
|---|---|
| G18V | -7.34 |
| A76V | -7.09 |
| **R206Q** | **-10.203** |
| N259S | -7.167 |
| **F284L** | **-9.822** |
| **Y371H** | **-10.308** |
| **F378I** | **-15.774** |
| **C384G** | **-15.719** |
| **D388Y** | **-12.682** |
| **C396R** | **-13.779** |
| **H398Y** | **-10.871** |
| **C404Y** | **-13.215** |
| **R420Q** | **-12.8** |
| R559Q | -4.044 |
| **R563C** | **-7.975** |
| T645C | -5.014 |
| A741V | -3.968 |

**Figure S1. In silico modelling predicts pathogenicity of CBL missense mutations that cluster around the linker region and RING finger domain**

**A)** Likelihood ratio matrix (LLR) of CBL amino acid substitutions, generated using the output of ESM-2. The log ratio between all possible residues (y-axis) and the wild-type at each position in the protein (x-axis) is plotted, where negative values (red) represent regions of high constraint for substitution or mutation, while positive values (blue) represent regions that are permissive to substitution. **B)** Location of individual CBL missense variants (n=17) identified in ZERO in CBL protein 3D structure, generated using ESMfold. Variants are colored by pathogenicity score, where red is “pathogenic” and purple is “not pathogenic”. Functional domains are colored according to the key. **C)** Table of CBL mutant pathogenicity scores calculated using ESM-2. Scores of less than -7.5 or greater than 7.5 are considered pathogenic (bolded).

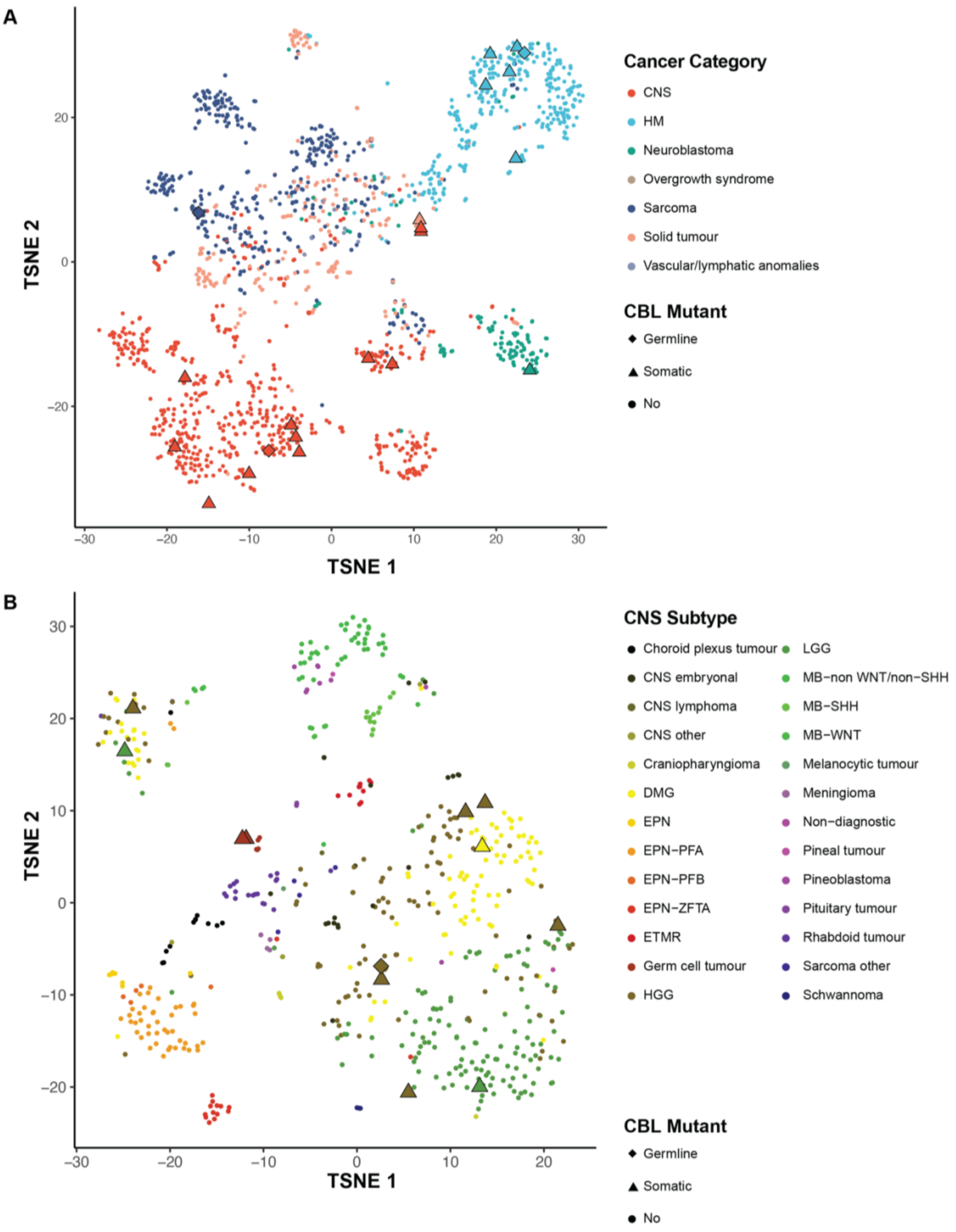


**Figure S2. CBL-mutated tumours have distinct gene expression profiles**

**A)** TSNE clustering of RNA sequencing data from all tumour types from the ZERO cohort (n=1653 samples) including samples with predicted functional CBL variants (n=21 samples). Cancer Category (CNS, hematological malignancy (HM), neuroblastoma, overgrowth syndrome, sarcoma, solid tumour, or vascular/lymphatic anomalies) and CBL mutant status are marked according to the key. **B)** TSNE clustering of RNA sequencing data from CNS tumours from the ZERO cohort (n=589 samples) including samples with predicted functional CBL variants (n=12 samples). CNS subtype and CBL mutation status are colored and marked according to the key.

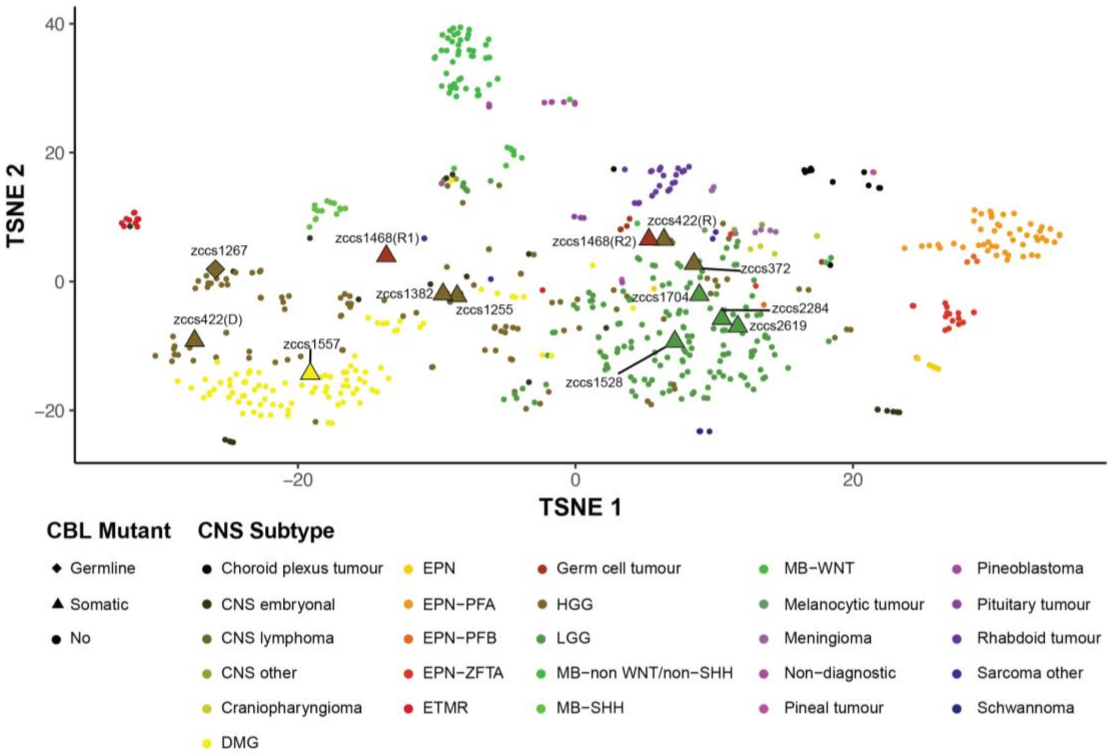


**Figure S3. CBL-mutated tumours have distinct DNA methylation profiles**

TSNE clustering of DNA methylation data from CNS tumours from the ZERO cohort (n=627 samples) including samples with predicted functional CBL variants (n=13 samples). DNA methylation data was unavailable for one sample. Diagnoses are colored and CBL mutated samples are marked according to the key. Patient-specific IDs of CBL mutated samples are marked.

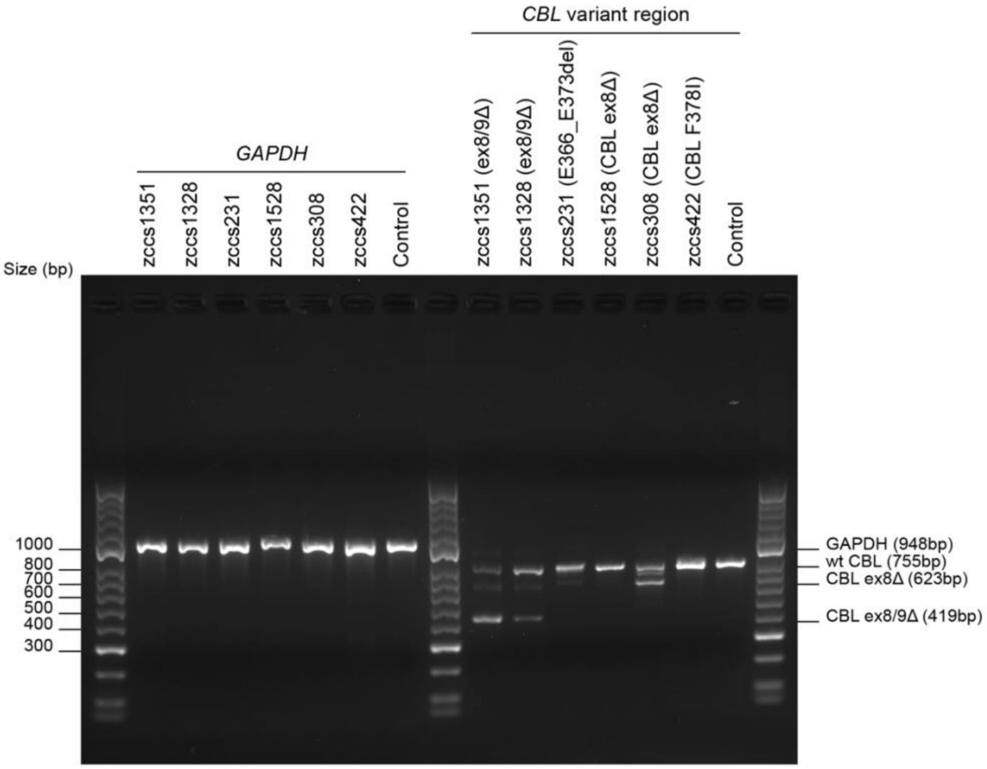


**Figure S4. PCR validation of *CBL* isoform expression in patient cDNA**

Primers targeting exons 6 and 11 were designed to amplify the CBL deletion/variant region from patient cDNA. Expected fragment sizes were as follows: 755bp (wt/ missense mutations), 623bp (ex8Δ) and 419bp (ex8/9Δ) and are marked on the right side of the gel image. GAPDH (948bp) was used as a control for cDNA quality and cDNA from a patient with wt CBL expression was used as a negative control for CBL alternate isoform expression.

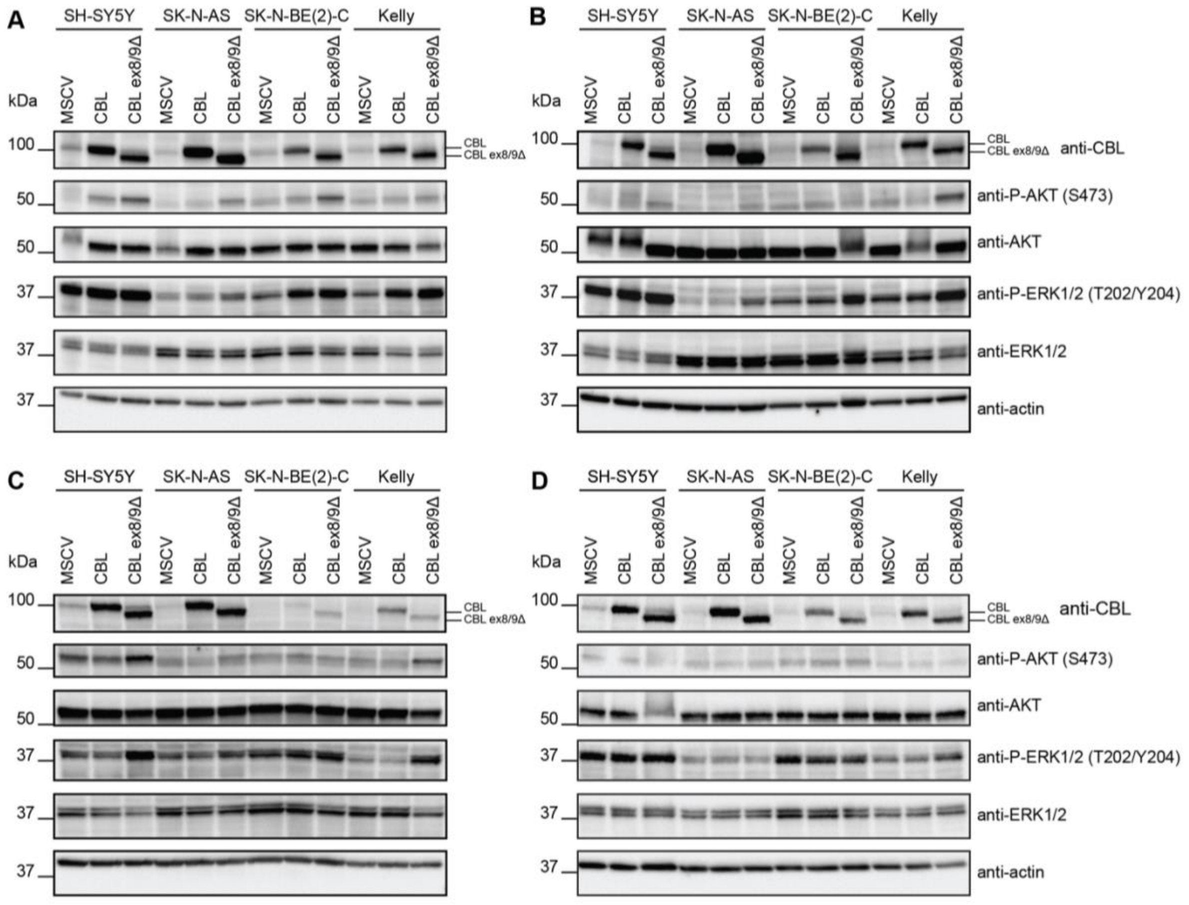


**Figure S5. Expression of CBL ex8/9Δ in neuroblastoma cell lines enhances intracellular signaling**

Western blot analysis of neuroblastoma cell lines (SH-SY5Y, SK-N-AS, SK-N-BE(2)-C and Kelly) transduced with CBL (wt), CBL ex8/9Δ or an empty vector control (MSCV). Cell lysates were harvested from four independent cell line passages (A-D).

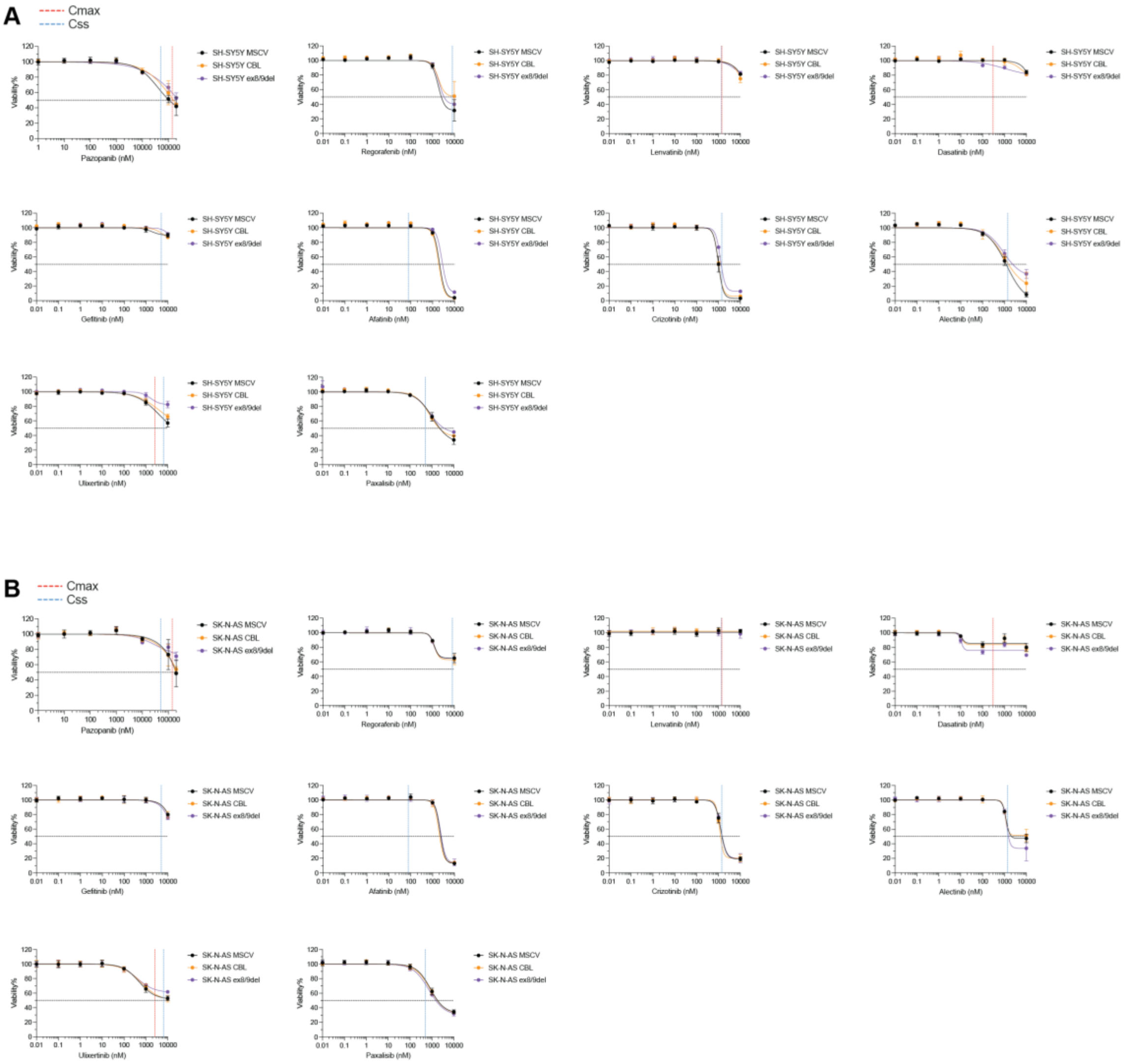


**Figure S6. High-throughput drug screening characterizes the impact of CBL variant expression on drug response in non-MYCN amplified neuroblastoma cell lines**

Dose response curves of non-MYCN amplified neuroblastoma cell lines, SH-SY5Y (A) and SK-N-AS (B), transduced with CBL variants and treated with a range of kinase inhibitor drugs: pazopanib, regorafenib, lenvatinib, dasatinib, gefitinib, afatinib, crizotinib, alectinib, ulixertinib, and paxalisib. Viability was determined using the CellTiter-Glo® 2.0 Cell Viability Assay and luminescence values were normalised to untreated and 100% death controls. Maximum serum concentration (Cmax) and steady-state concentration ($C_{ss}$) of each of the drugs, if known, is represented by the red and blue dotted lines, respectively. Starting viability for each cell line is depicted on the y-axis by the dotted lines. Cells were screened in technical triplicates in three independent experiments and data is presented as mean ± SEM (n=3).

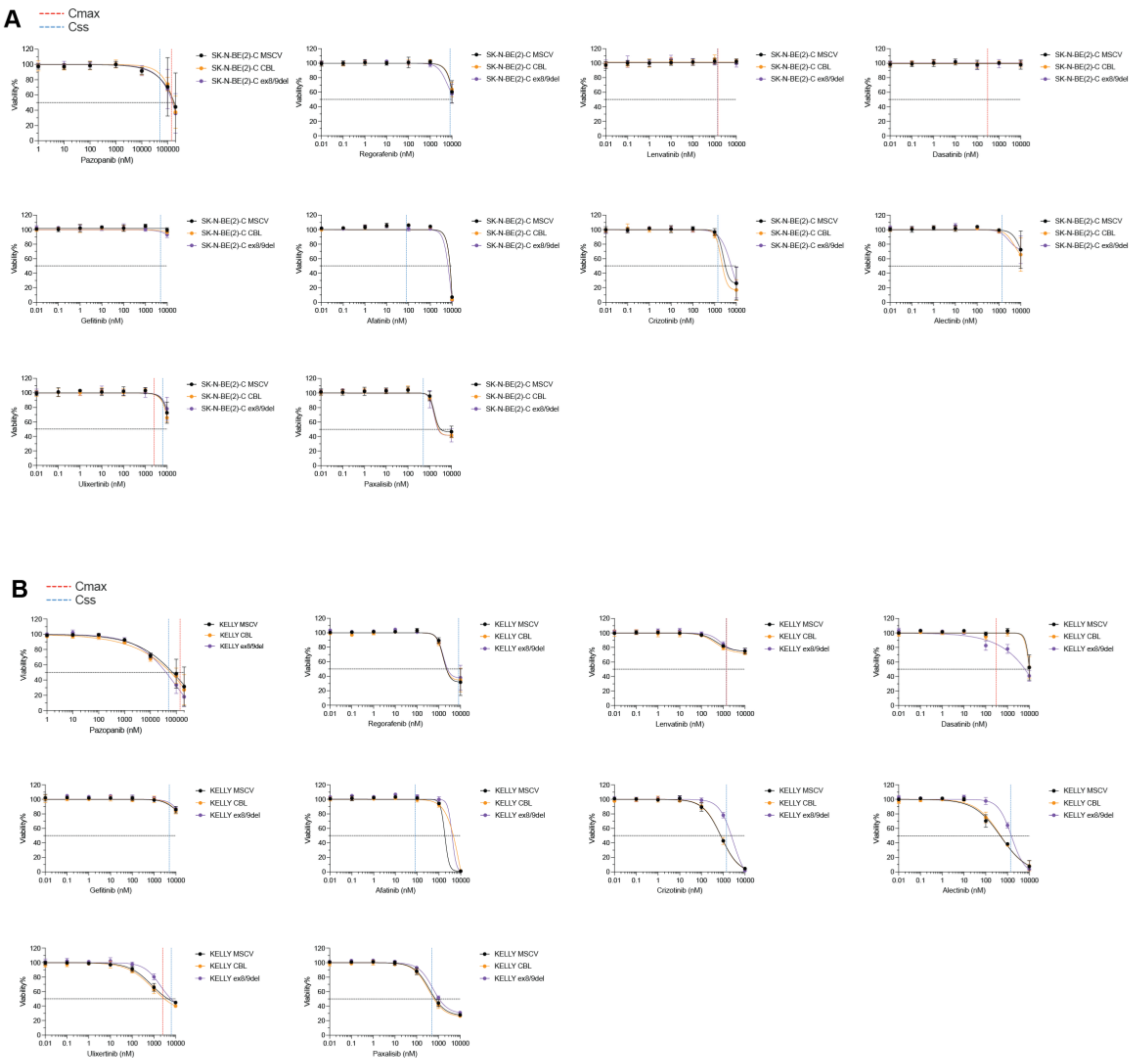


**Figure S7. High-throughput drug screening characterizes the impact of CBL variant expression on drug response in MYCN amplified neuroblastoma cell lines**

Dose response curves of MYCN amplified neuroblastoma cell lines, SK-N-BE(2)-C (A) and Kelly (B), transduced with CBL variants and treated with a range of kinase inhibitor drugs: pazopanib, regorafenib, lenvatinib, dasatinib, gefitinib, afatinib, crizotinib, alectinib, lorlatinib, trametinib, ulixertinib, and paxalisib. Lorlatinib and trametinib plots are also included in Fig. 3C of the main text. Viability was determined using the CellTiter-Glo® 2.0 Cell Viability Assay and luminescence values were normalised to untreated and 100% death controls. Maximum serum concentration (Cmax) and steady-state concentration ($C_{ss}$) of each of the drugs, if known, is represented by the red and blue dotted lines, respectively. Starting viability for each cell line is depicted on the y-axis by the dotted lines. Cells were screened in technical triplicates in three independent experiments and data is presented as mean ± SEM (n=3).

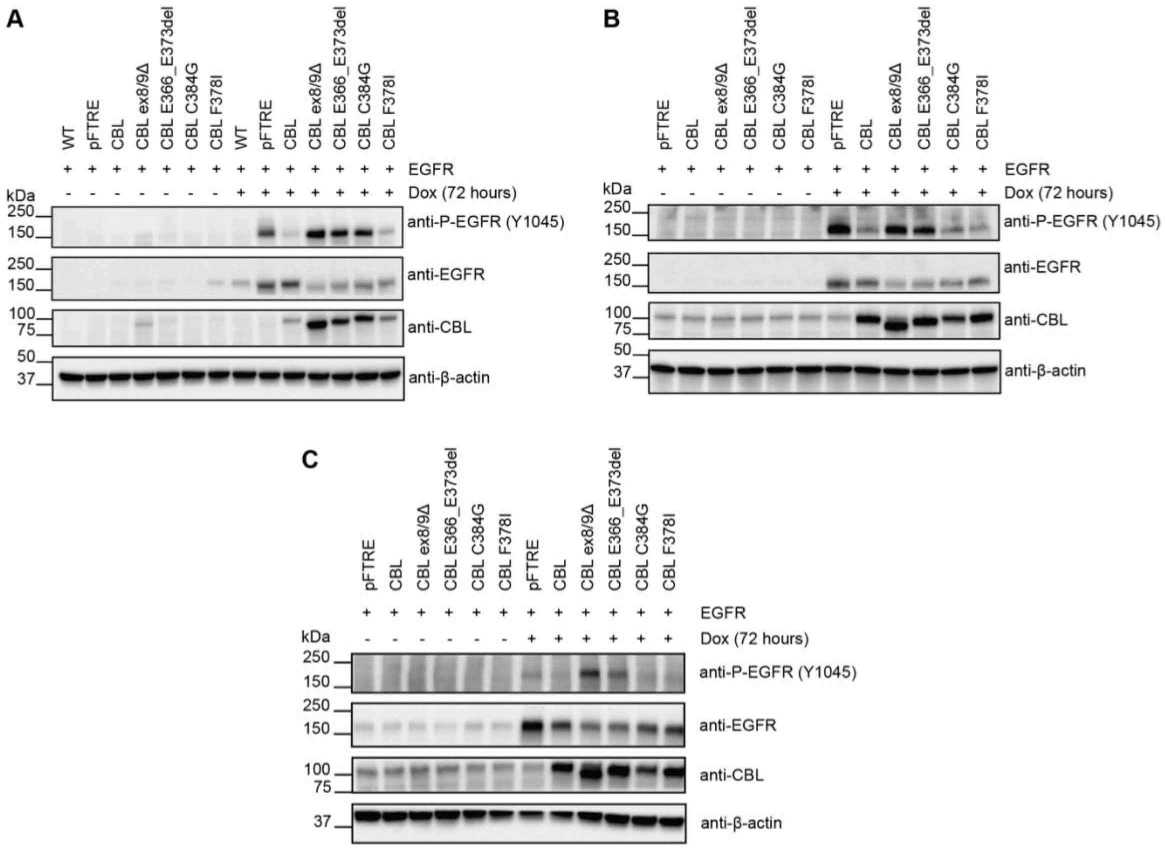


**Figure S8. Co-expression of EGFR with CBL E366_E373del, or CBL ex8/9Δ maintains phosphorylation of EGFR**

Western blot analysis of three biologically independent Ba/F3 cell lines (A-C) with doxycycline (dox) inducible expression of EGFR and CBL variants.
